# Structure of a Type II SMC Wadjet Complex from *Neobacillus vireti*

**DOI:** 10.1101/2025.03.10.642339

**Authors:** Florian Roisné-Hamelin, Hon Wing Liu, Stephan Gruber

## Abstract

Structural Maintenance of Chromosomes proteins are essential DNA-folding motors that facilitate critical cellular functions, including chromosome segregation and DNA repair. Wadjet systems are prokaryotic SMC complexes specialized in cellular immunity against plasmids. Type I Wadjet systems restrict plasmids via a DNA extrusion-cleavage reaction. Two other Wadjet types (II and III) have also been identified, however, their molecular characteristics are unclear. Here, we reconstituted a representative type II Wadjet system from *Neobacillus vireti*. We show that this system shares substrate selection and cleavage properties with type I but exhibits distinctive structural features, including a long elbow-distal coiled coil, a channel-less hinge, and a tandem KITE subunit. These features help identify the common and distinguishing architectural elements in the family of Wadjet systems and raise intriguing questions about the evolution of prokaryotic SMC complexes.

## Introduction

Structural Maintenance of Chromosome (SMC) complexes are ATP-driven motors that fold DNA through loop extrusion, a process in which they progressively enlarge DNA loops. SMC complexes facilitate essential cellular functions, including chromosome condensation and segregation (*e.g.* cohesin and condensin in eukaryotes and bacterial counterparts Smc-ScpAB, MukBEF, and MksBEF), DNA repair (Smc5/6 and cohesin), and cellular immunity against invasive DNA elements (Smc5/6 in eukaryotes and Wadjet in prokaryotes) ^1,2^.

SMC motor units include a dimer of SMC proteins with an ABC-type ATPase head domain from which a (roughly 50 nm) long antiparallel coiled coil extends and terminates in a dimerization component, the hinge. The kleisin protein connects the head-proximal coiled coil of one SMC (the v-SMC, at the neck interface) to the bottom of the ATPase head of the other SMC subunit (the κ-SMC, at the cap interface) and recruits a dimer of accessory KITE proteins (in prokaryotic SMC complexes and Smc5/6) or two HAWK subunits (in cohesin and condensin) (Figure 1A, displaying Wadjet as an example). KITE proteins harbour two tandem winged-helix domains (WHD) with the amino-terminal WHD mediating KITE protein dimerization. The KITE dimer binds to central kleisin sequences and contributes to DNA binding ^3^. Certain SMC complexes, such as bacterial MukBEF and Wadjet, form stable dimers, with two motor units held together by the dimerization of sequences found at the kleisin amino-terminus, namely an amino-terminal WHD (‘nWHD’) and an α-helical bundle ^4–6^.

**Figure 1:**
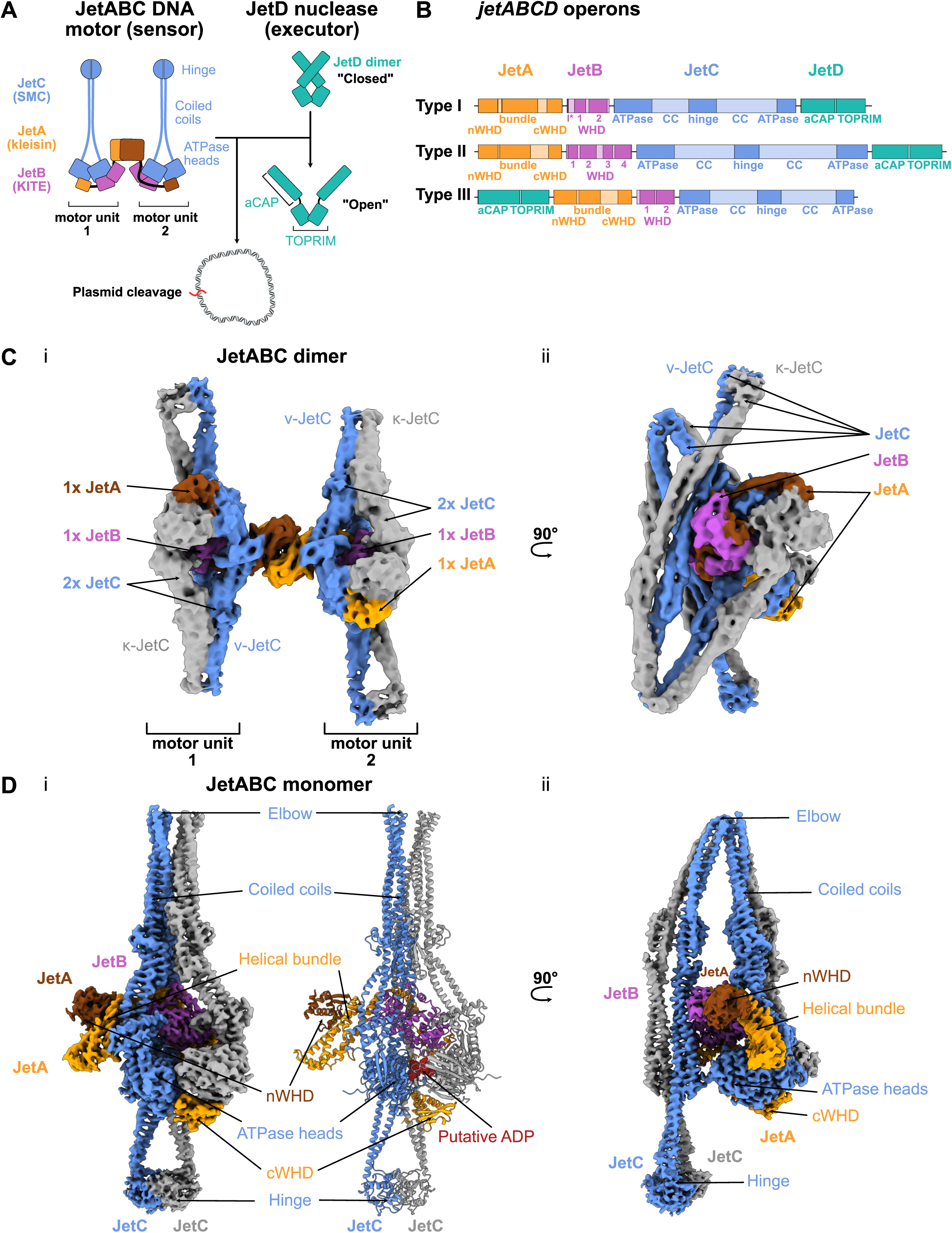
Cryo-EM structures of a type II Wadjet system from *Neobacillus vireti*. (A) Schematic representation of the *E. coli* GF4-3 Wadjet-I ecJetABC dimer in the resting state ^5^. (B) Schematic representation (drawn to scale) of representative operon of type I (*E. coli GF4-3*), type II (*N. vireti*) and type III (*B. thuringiensis*) ^5^. Protein domains have been identified using available related protein structures or AlphaFold2 predictions as a guide ^5,28^. Amino-terminal WHD, nWHD; carboxy-terminal WHD, cWHD, bundle: α-helical bundle; ATPase: ATPase cassette; CC: coiled coils. I*: type I JetB IFDR motif binding to JetD aCAP (C) Structure of nvJetABC dimer (overall resolution: 6.71 Å, PDB: 9QE0) showing the global architecture of the nvJetABC sub-complex, with JetA subunits in orange and brown colours, JetB in violet, v-JetC (responsible of the neck interface) in blue, k-SMC (responsible of the cap interface) in grey. (Ci) face view, (Cii) side view of the density map (90°-rotated along the y axis). (D) Structure of a nvJetABC monomer (overall resolution: 3.46 Å, PDB: 9QE1) with the corresponding model, showing the organization of the nvJET-II motor unit (same colors as in (C)). (i) and (ii): face and side views, respectively. See also Table 1, Figure S1, S2, S3, and S4.

**Table 1.**
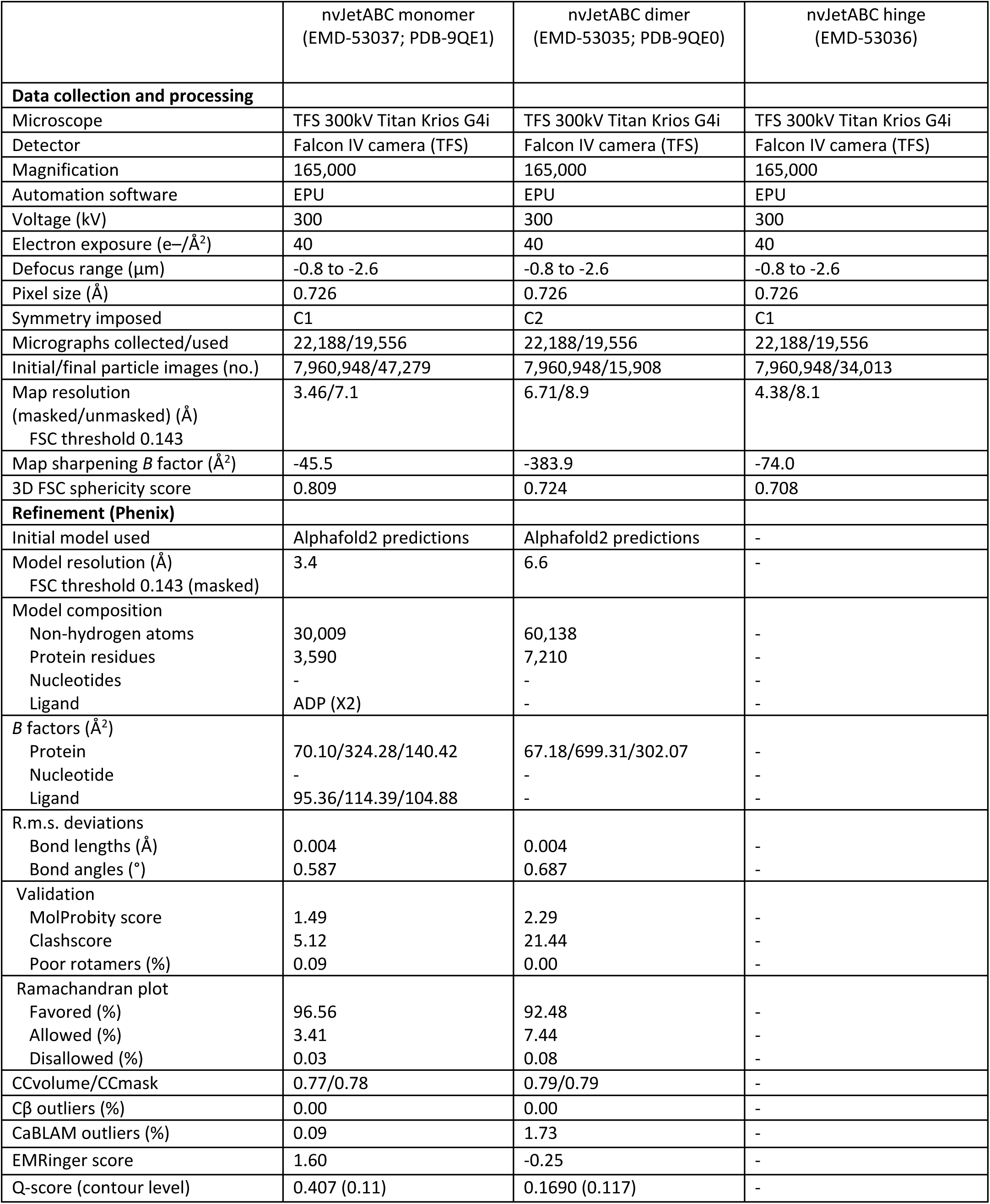
Cryo-EM data collection, processing, and model building. Related to Figures 1, 2, 3.

|  | nvJetABC monomer<br>(EMD-53037; PDB-9QE1) | nvJetABC dimer<br>(EMD-53035; PDB-9QE0) | nvJetABC hinge<br>(EMD-53036) |
| --- | --- | --- | --- |
| **Data collection and processing** |  |  |  |
| Microscope | TFS 300kV Titan Krios G4i | TFS 300kV Titan Krios G4i | TFS 300kV Titan Krios G4i |
| Detector | Falcon IV camera (TFS) | Falcon IV camera (TFS) | Falcon IV camera (TFS) |
| Magnification | 165,000 | 165,000 | 165,000 |
| Automation software | EPU | EPU | EPU |
| Voltage (kV) | 300 | 300 | 300 |
| Electron exposure (e<sup>-</sup>/Å<sup>2</sup>) | 40 | 40 | 40 |
| Defocus range (μm) | -0.8 to -2.6 | -0.8 to -2.6 | -0.8 to -2.6 |
| Pixel size (Å) | 0.726 | 0.726 | 0.726 |
| Symmetry imposed | C1 | C2 | C1 |
| Micrographs collected/used | 22,188/19,556 | 22,188/19,556 | 22,188/19,556 |
| Initial/final particle images (no.) | 7,960,948/47,279 | 7,960,948/15,908 | 7,960,948/34,013 |
| Map resolution<br>(masked/unmasked) (Å)<br>FSC threshold 0.143 | 3.46/7.1 | 6.71/8.9 | 4.38/8.1 |
| Map sharpening *B* factor (Å<sup>2</sup>) | -45.5 | -383.9 | -74.0 |
| 3D FSC sphericity score | 0.809 | 0.724 | 0.708 |
| **Refinement (Phenix)** |  |  |  |
| Initial model used | AlphaFold2 predictions | AlphaFold2 predictions | - |
| Model resolution (Å)<br>FSC threshold 0.143 (masked) | 3.4 | 6.6 | - |
| Model composition |  |  |  |
| Non-hydrogen atoms | 30,009 | 60,138 | - |
| Protein residues | 3,590 | 7,210 | - |
| Nucleotides | - | - | - |
| Ligand | ADP (X2) | - | - |
| *B* factors (Å<sup>2</sup>) |  |  |  |
| Protein | 70.10/324.28/140.42 | 67.18/699.31/302.07 | - |
| Nucleotide | - | - | - |
| Ligand | 95.36/114.39/104.88 | - | - |
| R.m.s. deviations |  |  |  |
| Bond lengths (Å) | 0.004 | 0.004 | - |
| Bond angles (°) | 0.587 | 0.687 | - |
| Validation |  |  |  |
| MolProbity score | 1.49 | 2.29 | - |
| Clashscore | 5.12 | 21.44 | - |
| Poor rotamers (%) | 0.09 | 0.00 | - |
| Ramachandran plot |  |  |  |
| Favored (%) | 96.56 | 92.48 | - |
| Allowed (%) | 3.41 | 7.44 | - |
| Disallowed (%) | 0.03 | 0.08 | - |
| CCvolume/CCmask | 0.77/0.78 | 0.79/0.79 | - |
| Cβ outliers (%) | 0.00 | 0.00 | - |
| CaBLAM outliers (%) | 0.09 | 1.73 | - |
| EMRinger score | 1.60 | -0.25 | - |
| Q-score (contour level) | 0.407 (0.11) | 0.1690 (0.117) | - |

ATP binding and hydrolysis mediate engagement and disengagement of the SMC head domains, respectively, and concomitant opening and closure of the head-proximal SMC coiled coils, which is thought to drive the process of DNA loop extrusion ^2,7–9^. While loop extrusion has been directly observed in single-molecule assays for various complexes ^10–14^, the molecular mechanism underlying this process remain incompletely understood ^9,15–18^.

Most prokaryotes rely on Smc-ScpAB for chromosome segregation ^19^. However, some branches of γ-proteobacteria have apparently lost Smc-ScpAB and instead utilize MukBEF for chromosome segregation ^19,20^. MukBEF promotes long-range DNA contacts throughout the chromosome, presumably through DNA loop extrusion ^21,22^. A third bacterial SMC complex, MksBEF, has been identified scattered across the phylogenetic tree and shares distant sequence homology with MukBEF ^23,24^. MksBEF promotes shorter long-range DNA contacts in *Pseudomonas aeruginosa* and was shown to serve as a backup segregation system when Smc-ScpAB activity is compromised ^22^. The evolutionary origins of MukBEF and MksBEF are possibly linked to Wadjet DNA restriction complexes ^1,25^.

Wadjet (also known as JetABCD, the JET nuclease, MksBEFG, or EptABCD) is a prokaryotic SMC complex involved in cellular immunity against invasive DNA elements which restricts circular plasmids ^1,4,5,24,26,27^. Three types of Wadjet systems (I-III) have been identified in their operon organization and domain composition ^26^ (Figure 1B). Wadjets are present in approximately 5 % of bacterial species ^24,26^ and comprise a SMC DNA sensor (JetABC) coupled with a TOPRIM domain-containing nuclease effector, JetD. Purified type I Wadjet (Wadjet-I) systems cleave DNA circles at random positions, independent of sequence or DNA helical topology, while leaving linear molecules intact ^4,5,28^. Notably, Wadjet-I struggles to bypass micrometre-scaled obstacles attached to the plasmid DNA ^28^, indicating that Wadjet motors survey DNA circular topology through their DNA loop extrusion activity. This notion aligns with a recent single-molecule imaging study ^13^.

Cryogenic electron microscopy (Cryo-EM) with single particle analysis revealed that the Wadjet-I DNA sensor is a dimer of SMC motor units ^4,5,28^, while JetD forms a dimer whose structure was determined in open ^29^ or closed, presumably inactive, conformations where the catalytic residues of the TOPRIM domains are shielded from access to DNA by adjacent arm-CAP (aCAP) domains ^4^. Plasmid-bound JetABC-I structures revealed a drastic conformational switch of the SMC motor units compared to the resting state. In the cleavage state, the motor units entrap a U-shaped DNA segment, which appears to be converted into a kinked V-shaped cleavage substrate upon JetD nuclease binding ^28^. Current understanding of Wadjet systems is primarily based on type I systems, whereas the other two types (Wadjet-II and Wadjet-III) remain largely uncharacterized ^4,5,13,28,29^. Generally, bacterial defence systems exist in many variations, which is likely due to the presence of phage counter-defence mechanism, driving rapid evolutionary diversification^30^. The implications of different Wadjet sequences and architectures on plasmid restriction activity and loop extrusion properties remain unknown.

Here, we have reconstituted a Wadjet-II system from *Neobacillus vireti* strain LMG21834 (nvJET-II) and characterized its plasmid cleavage properties. We uncover commonalities with Wadjet-I but also differences. nvJET-II demonstrates DNA cleavage only in the presence of manganese, otherwise exhibiting biochemical characteristics analogous to a previously described Wadjet-I system from *E. coli* (ecJET-I)^5^. Specifically, nvJET-II cleaves circular (but not linear) DNA at random positions through an ATP- and JetABCD-dependent manner. nvJET-II cleaves linear DNA that has been artificially circularized, suggesting that it is a DNA shape-sensing system like ecJET-I. Cryo-EM revealed distinctive features of the nvJET-II motor—its SMC subunits display a specific coiled coil arrangement, with a hinge dimerization interface lacking a central channel, thus more closely resembling MukBEF. Intriguingly, nvJET-II harbors a tandem KITE protein with four tandem WHDs, which is the functional equivalent of a KITE dimer found in other SMC complexes. This unique organization provides an opportunity to evaluate the contributions of individual KITE WHDs during loop extrusion.

## Results

### Reconstitution of a Wadjet-II system from *Neobacillus vireti*

To understand how different types of Wadjet vary from one another and from the previously characterized Wadjet-I systems, we selected a Wadjet-II system (nvJET-II) from *Neobacillus vireti* strain LMG 21834 for in-depth characterization ^5,26^. This system, when introduced to *Bacillus subtilis*, robustly restricts plasmids regardless of replication origin and propagation mode ^5,26^. Notably, like most or all other Wadjet-II systems, nvJET-II exhibits significantly larger SMC and KITE genes (Figure 1B) ^26^. nvJetABC and nvJetD proteins were recombinantly expressed in *E. coli* and purified (Figure S1A, Methods). nvJetABC harboured ATPase activity at a rate of approximately 100 ATP molecules per complex per minute at 37°C, with no noticeable stimulation by nvJetD (Figure S1B). This is about fourfold lower when compared to the ATPase activity of ecJET-I under the same conditions ^5^ but much higher than the rate reported for a closely related Wadjet-I system from a *P. aeruginosa* strain PA14 ^4^, implying that these differences are type-unrelated. DNA presence slightly enhanced ATPase activity for nvJET-II (Figure S1B), akin to ecJET-I^5^ and other SMC complexes ^31,32^.

### Cryo-EM analysis of nvJetABC

We investigated the structure of nvJetABC using cryo-electron microscopy. To this end, nvJetABC complexes freshly eluted from an analytical size exclusion chromatography column were pre-incubated with ATP and frozen (see Methods). We determined two structures: a holo-complex of nvJetABC with two SMC motor units at a lower resolution (overall: 6.8 Å, Figure 1C) and a single nvJetABC motor unit at 3.4 Å estimated overall resolution (Figure 1D, see also Figures S2, S3, S4 and Table 1), both in the absence of DNA in a putative “resting” conformation. nvJetABC motor units harbour four subunits with a JetA_1_B_1_C_2_ stoichiometry. The SMC motor units dimerize primarily through the nWHD of kleisin JetA making the dimer-of-tetramers holo-complex (Figure 1C). Notably, the overall geometry of nvJET-II more closely resembles the MukBEF and diverges from the V-shaped ecJET-I resting state geometry (Figure 2A). The two SMC motor units are positioned side-by-side in a plane but are rotated by almost 180° relative to each other. To assess whether this nvJET-II dimer geometry might be representative of the Wadjet-II family, we predicted the structures of several Wadjet-II SMC motors from a diverse range of species using AlphaFold3 ^33^ showing similar overall architectures (Figure S5A), suggesting that nvJET-II is indeed a representative Wadjet-II member (or a bias in AF3). Together, these findings indicate that the V-shaped resting state of ecJET-I might be a structural peculiarity of Wadjet-I systems, but the functional role of this specific geometry is unclear. As observed for the ecJET-I resting state, the heads display an ATP-nonengaged state, with electron density resembling a nucleotide (possibly ADP) being present at the nvJetC heads (Figure 1D, Figure S4D).

**Figure 2:**
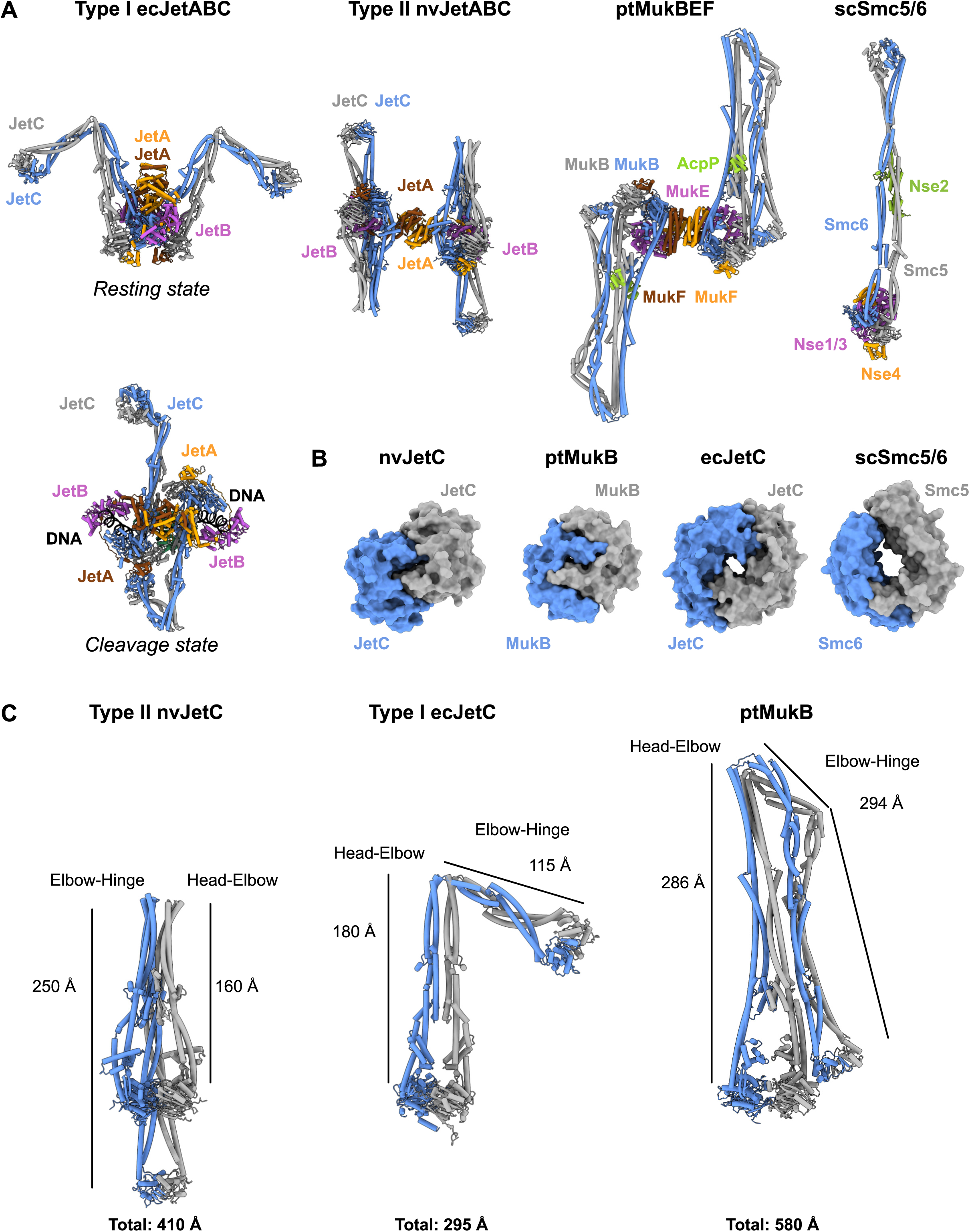
Comparison of nvJET-II with other SMC complexes. (A) Comparison of the model of nvJET-II (JetABC dimer, PDB: 9QE0) with ecJET-I reconstructions ^5,28^, ptMukBEF dimer (PDB: 7NZ4), scSmc5/6 (PDB: 7QCD)^6,50^. The ecJET-I resting and cleavage competent state reconstructions were made by extending the deposited models (respectively PDB: 8BFN and PDB: 8Q72) at the upper part of the coiled coils using AlphaFold2 predictions. Same coloring as in Figure 1. (B) Comparison of hinge shapes of nvJetC and ecJetC (AlphaFold2 predictions), ptMukB (PDB: 7NZ4) and scSmc5/6 (PDB: 7QCD). Only the hinge domain is displayed, with SMC chains displayed in blue and grey colors, respectively. (C) Comparison of the coiled coil length for nvJET-II (PDB: 9QE1), ecJET-I (PDB: 8BFN-based reconstruction), and ptMukBEF (PDB: 7NZ4). See also Figure S5 and S6.

The density corresponding to the nvJetC hinge showed lower resolution, but we nevertheless obtained a map of the hinge region using local 3D classification followed by local refinement (Figure S2, S3C). An AlphaFold2 model of the nvJetC hinge fits well into this cryo-EM density (Figure S4E) ^34^. While the predicted structure of the ecJetC hinge displays a “donut-like” fold, consistent with typical SMC hinge structures ^35^, the prediction as well as the experimental structure of the nvJetC hinge more closely resemble the one of MukB, characterized by a more compact and rectangular shape lacking a central channel (Figure 2B, Figure S4E, S5B).

nvJetC includes an elbow, a prominent folding point in the coiled coil, as in MukBEF and ecJET-I complexes ^4–6,36^. Intriguingly, in nvJET-II, the head-to-elbow coiled coil is significantly shorter than the elbow-to-hinge coiled coil, thus positioning the hinge and elbow on opposite sides of the nvJetC head module (Figure 1C, 1D, 2C). The elbow-to-hinge coiled coil directly contacts the nvJetC ATPase head as well as the nvJetB KITE, with the latter being sandwiched between the nvJetC coiled coil and the heads (Figure 1C). The position of the KITE protein on top of the SMC heads is comparable with MukBEF, Smc5/6, and the ecJET-I cleavage state, yet again different from ecJET-I in the resting state (Figure 2A, Figure S6A).

JetD was not present in our cryo-EM preparations. However, we conducted structural comparisons using AlphaFold3 predictions (Figure S5A ii, S6B, S6C) ^33^. Previously obtained structures of JetD-I ^4,29^ revealed a protein with two main components: a TOPRIM domain (likely containing the cleavage catalytic site) and a flexibly connected aCAP (a CAP domain with an ‘arm’ extension), which exhibits limited similarity with topoisomerase CAP domains, but lacks the conserved catalytic tyrosine ^4,5^. The predicted nvJetD-II dimer also contains TOPRIM and CAP-like domains and shares an analogous dimer organization (Figure S6B ii). While the nvJetD TOPRIM domain largely superimposes with ecJetD (with local variations), its aCAP differs significantly, particularly in the amino-terminal sequences that is known to connect JetD-I with JetABC-I (Figure S6C iii) ^4,28^. These predicted structural differences suggest that nvJetD-II interacts with nvJetABC-II distinctly from Wadjet-I systems, indicating significant evolutionary diversification of type I and II JetD proteins.

### Wadjet-II complexes contain a tandem KITE subunit

Another distinguishing feature of nvJET-II concerns its KITE protein. KITE-containing SMC complexes typically feature a dimer of KITE proteins per motor unit—homodimers in prokaryotic SMC complexes and heterodimers in eukaryotic Smc5/6 ^3^. These dimers form through homo- or heterotypic interactions of the amino-terminal WHD of the respective KITE proteins. nvJET-II has a single tandem KITE subunit per motor unit, comprising four tandem WHDs (denoted WHD-1 to WHD-4) (Figure 1, Figure 3). This tandem KITE protein structurally resembles a typical KITE dimer found in other prokaryotic SMC complexes, similarly spanning the central sequences of the kleisin nvJetA (Figure 3A, 3B). Each half of tandem KITE protein (here termed a ‘lobe’) corresponds to an individual KITE protein in other SMC motor units (Figure 3B). Since most or all Wadjet-II KITE genes are larger (388 residues in nvJetB) when compared to other KITE genes (242 residues in ecJetB), they might all represent tandem KITE proteins (Figure S5A) ^26^. Structure-based sequence alignment using FoldMason indicates a substantial divergence between the two nvJetB lobes (Figure 3C) ^37^.

**Figure 3:**
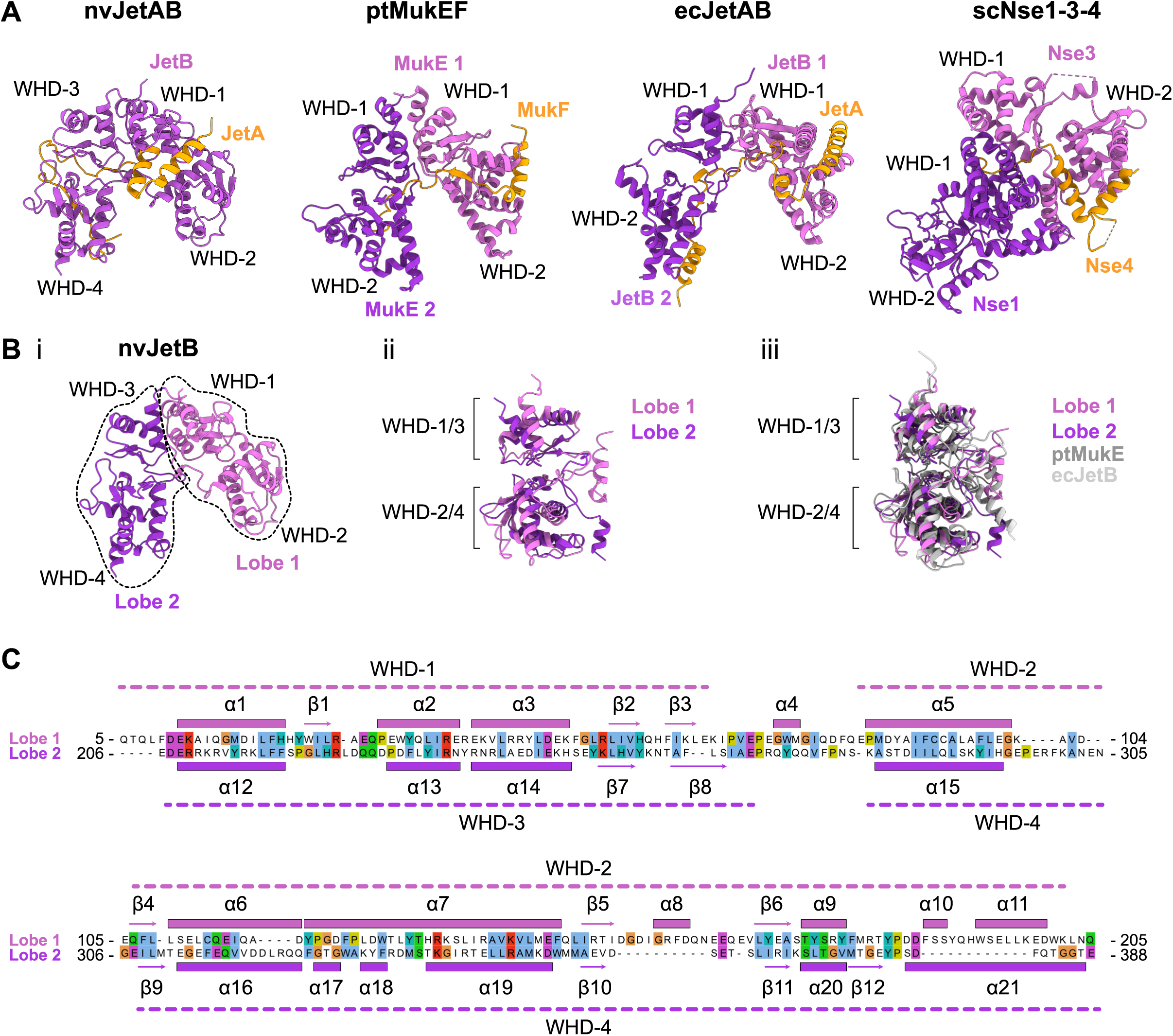
The nvJET-II tandem KITE. (A) Structural comparison of the nvJetB tandem KITE protein (PDB: 9QE1) with other SMC KITE dimers (ptMukE, PDB: 7NZ4; ecJetB, PDB: 8AS8; scNse1/3, PDB: 7QCD). The KITE proteins are shown in shades of pink and violet, with the central part of the kleisin bound to the KITES displayed in orange. (B) The two lobes of the nvJetB tandem KITE protein (i). (ii) Superimposition of nvJetB lobe 1 (nvJetB 5-205) with lobe 2 (nvJetB 206-388). (ii). Structural alignment of nvJetB lobes with ptMukE (PDB: 7NZ4) and ecJetB (PDB: 8AS8) using FoldMason ^37^. (C) Structure-based sequence alignment of nvJetB lobe 1 (5-205) and lobe 2 (206-388) using FoldMason ^37^. The position of α-helices (α1-α21, displayed as squares) and β-sheets (β1-12, displayed as arrows) is shown for lobes 1 and 2 (in pink and violet, respectively). The positions of the WHD domains (WHD-1 to WHD-4) are shown in dashed lines. See also Figure S5 and S6.

The tandem protein might have emerged by KITE gene duplication and fusion, followed by sequence diversification. Consistent with this evolutionary model, WHD-1 and WHD-3 interact at the center of the tandem KITE protein. WHD-2 and WHD-1 from the first lobe bind to more amino-terminal and WHD-3 and WHD-4 from the second lobe to more carboxy-terminal sequences of kleisin JetA. Moreover, a longer linker connects WHD-2 in the first lobe to WHD-3 in the second lobe. KITE dimers mediate DNA binding in at least three states ^6,25,28^. The Wadjet-II tandem-KITE likely plays equivalent roles. This unique tandem KITE characteristic within nvJET-II systems will offer an opportunity to investigate the contributions of the four WHDs individually in the process of DNA loop extrusion in a prokaryotic SMC complex.

### DNA cleavage properties of nvJET-II

With the available structures displaying similarities and differences between Wadjet-I and Wadjet-II complexes, we next wondered whether they share related DNA cleavage activities (Figure 4, Figure S7). When incubated in standard Wadjet-I DNA cleavage buffer (*i.e.* ATG buffer: Hepes pH 7.5 10 mM, KOAc 150 mM, MgCl_2_ 5 mM; Liu et al., 2022), nvJET-II proteins showed minimal (if any) DNA cleavage activity. Addition of manganese chloride enabled plasmid cleavage by nvJET-II, with optimal activity observed at a concentration of 4-5 mM MnCl_2_ (Figure 4A, S7A, S7B). In contrast, ecJET-I cleaved plasmid DNA regardless of manganese presence. Plasmid cleavage by nvJET-II required both nvJetABC and nvJetD proteins, as well as ATP (Figure 4B). Pre-linearized DNA remained unaltered by nvJET-II (Figure 4C). together suggesting active DNA scanning by SMC motors prior to cleavage as previously proposed for ecJET-I ^28^.

**Figure 4:**
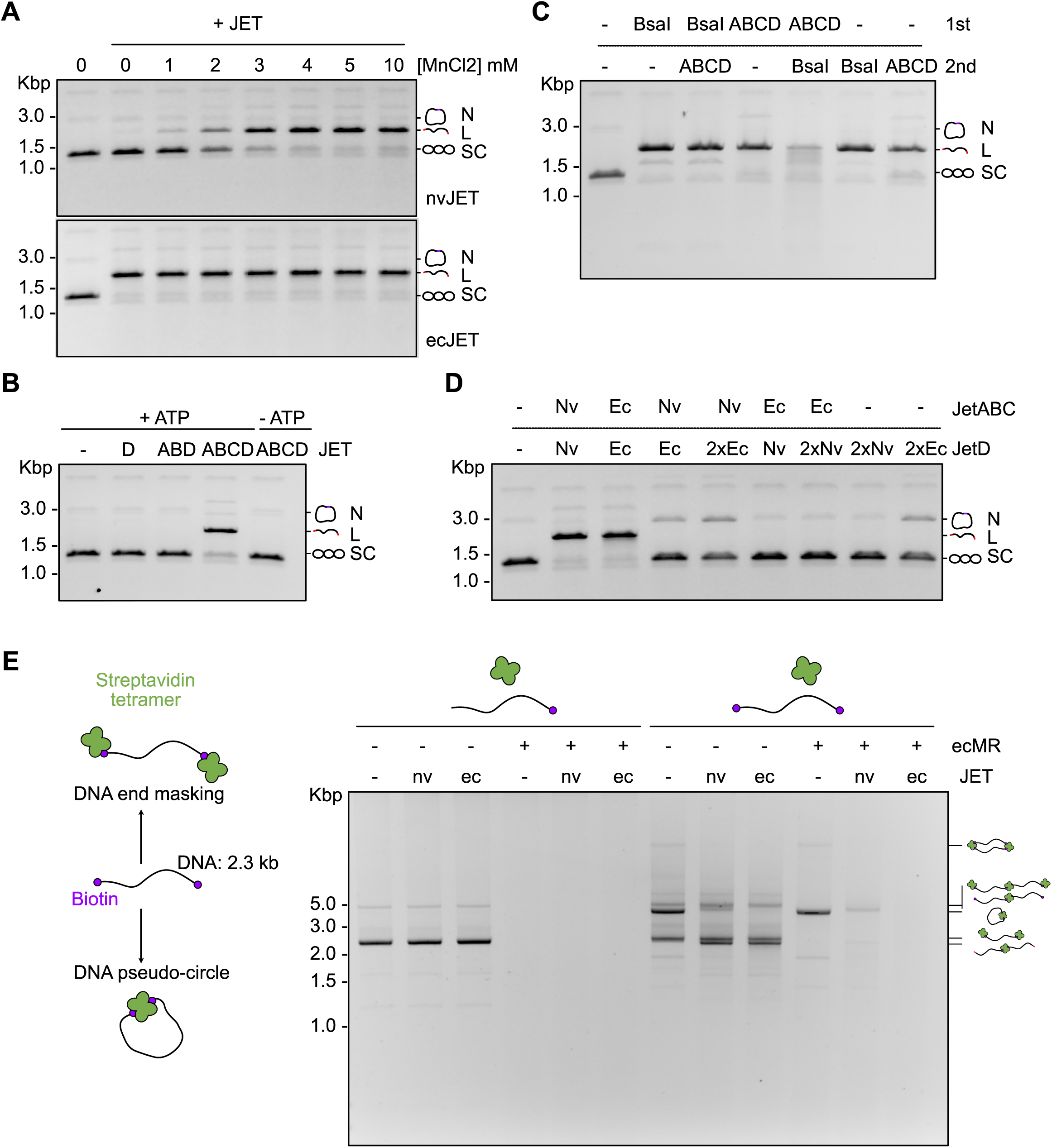
Biochemical characterization of nvJET-II plasmid cleavage activity. (A) Plasmid DNA cleavage reactions with ecJetABCD (12.5 nM JetABC dimer, 25 nM JetD dimer) or nvJetABCD (25 nM JetABC dimers, 50 nM JetD dimer) incubated with pDonor (8.75 nM) 10 min at 37°C in presence of ATP (1 mM) and MgCl2 (as indicated). The reaction was stopped on ice, followed by SDS/heat treatment, and the products were analysed by ethidium bromide agarose gel electrophoresis. The resulting DNA species are labelled: N for Nicked; L for Linear; SC for supercoiled. The same labelling is used in the following panels. (B) Plasmid cleavage assay with nvJET-II subcomplexes (nvJetABC, nvJetD, or nvJetABCD) in presence or absence of ATP (1 mM). (C) Post-treatment (10 min, 37°C) of nvJET-II cleaved plasmid DNA by the restriction enzyme BsaI resulted in a DNA smear. (D) Plasmid cleavage assay with a mixing of ecJET-I and nvJET-II subcomplexes. (E) Binding of streptavidin tetramers to double-biotinylated DNA molecules creates a heterogeneous mixture of species, including artificial pseudo-circular DNA circles and linear multimers ^28^. In contrast, streptavidin binding to single biotinylated DNA exclusively generates linear molecules. Left: schematic representation of streptavidin-mediated pseudo circularization of biotinylated DNA. Right: cleavage assay with ecJET-I (ecJetABC: 12.5 nM dimer, ecJetD: 25 nM dimer) or nvJET-II (nvJetABC: 25 nM dimer, nvJetD: 50 nM dimer), ecMR nuclease (125 nM, heterotetramer) on DNA species obtained by streptavidin (100 nM monomer) incubation with single or double biotinylated 2.3 kb DNA substrates (7 nM). The experiments shown are representative examples of at least two independent replicates. See also Figure S7 and S8.

As previously reported for two closely related Wadjet-I systems ^4,5^, we found that post-treatment of nvJET-cleaved DNA with a single cutter restriction enzyme resulted in DNA smearing, indicating that nvJET-II cleavage occurred at random positions (Figure 4C). Sequencing of nvJET-cleaved DNA fragments (after ligation) revealed widespread, sequence-nonspecific distribution of cleavage sites across the plasmid (Figure S7C). Examination of the generated DNA ends revealed 5’ overhangs of varying lengths (1-4 nt), a signature of ‘messy’ cleavage events by the nuclease nvJetD-II, as previously observed with ecJetD-I (Table S1) ^4,28^. As with Wadjet-I systems, nvJET-II cleavage products were sensitive to RecBCD exonuclease, indicating that nvJetD does not remain stably associated at the cleaved DNA ends (Figure S7D). Moreover, nvJET-II demonstrated the ability to cleave nicked plasmid DNA (Figure S7E), as well as plasmids of different sizes (up to 26.7 kb) (Figure S7F)*. In vivo*, DNA restriction activity by nvJET-II appears to wane with target DNA circle sizes of 50 kb and beyond, possibly indicating a similar, if not slightly more constricted size limit when compared to other types (Figure S8) ^5^.

nvJetD did not cleave DNA in the presence of ecJetABC, nor did ecJetD cleave DNA in the presence of nvJetABC (Figure 4D). These findings indicate that JetABC and JetD from different types are not interchangeable and suggest JetD activation relies on specific protein-protein interactions, as previously found for ecJET-I ^28^, rather than solely on DNA deformation by JetABC, potentially preventing accidental JetD activation on the chromosome. Finally, we examined nvJET-II’s ability to cleave (biotinylated) linear DNA artificially circularized by streptavidin addition (Figure 4E) ^28^. nvJET-II selectively cleaves pseudo-circular DNA species while leaving linear molecules intact, as previously observed with ecJET-I (Figure 4E). Co-treatment of DNA with nv/ecJET and linear DNA-targeting endonuclease ecMR (Mre11-Rad50; SbcCD) resulted in nearly complete elimination of all species (Figure 4E) ^28^. These findings indicate that although structurally distinct (at least in their resting states), nvJET-II senses the circular nature of DNA substrates prior to cleavage, in a manner that is remarkably similar to ecJET-I and suggests that all Wadjets discriminate plasmids via a common principle based on the sensing of DNA size and shape by an extrusion-cleavage mechanism. We suspect that key differences will emerge in the presence of native plasmid substrates which bring anti-Wadjet defense mechanisms into play targeting one or two Wadjet types but not all three.

## Discussion

Wadjet-I systems discriminate self-from nonself-DNA by measuring DNA size. Due to the reliance on complete DNA extrusion for DNA sizing, Wadjet-I systems are limited to recognizing circular DNA molecules, with all linear DNA (big or small) escaping Wadjet-I detection. ^4,5,13,28^. Our findings on nvJET-II indicate that Wadjet-II, and by extension likely also Wadjet-III, follows the same principle, implying that the extrusion-cleavage mechanism is conserved across all types of Wadjet systems. Whether the same holds true for Smc5/6 in viral restriction remains to be seen ^1^.

This study demonstrates that nvJET-II shares structural features with MukBEF and Wadjet-I but also has distinct characteristics. The similarities include a dimeric nature of the SMC complexes (Figure 1), with the individual core motor unit exhibiting a typical ring-shaped, extended SMC architecture resembling that of Wadjet-I and other SMC complexes (Figure 1C, 1D, 2A). These similarities suggest shared DNA motor mechanisms and an extrusion-cleavage mechanism likely involving a cleavage state with motor units holding bent DNA as in ecJET-I ^4,5,13,28^. The differences include an altered dimer geometry of the resting state, a unique aCAP fold of the nuclease subunit, a tandem KITE subunit, and a manganese requirement for activity. We suspect that these structural differences and the divergent protein sequence emerged due to evolutionary pressures from biological conflicts, *i.e.* the continuous need to escape plasmid anti-Wadjet mechanisms.

### Specific features of nvJET-II systems and implications for the plasmid restriction mechanism

The resting state structure of nvJET-II (Figure 1C, 1D) adopts a dimer geometry found in the MukBEF dimer ^6^, and differs from the ‘collapsed’ geometry of ecJET-I dimer ^5^, particular with regards to how the two SMC motor units connect to each other through the nWHD of kleisin (Figure 2A). Given these similarities in the resting states, we hypothesize that initial DNA loading events of nvJET-II may mirror those of MukBEF, involving the opening of the kleisin-N gate and initial non-topological DNA contacts by the KITE dimer recently shown for MukBEF ^25^.

When compared to MukB and ecJetC, nvJetC displays intermediary structural features. For example, the coiled coils are significantly longer (≃410 Å) than in ecJetC (≃295 Å) but shorter than in MukB (≃580 Å) (Figure 2C). This may relate to the capacity of obstacle threading through the SMC lumen. Whether the coiled coils must fully open during loop extrusion or only to the elbow position (especially when facing an obstacle), and how this affects loop extrusion properties, remain important questions with implications for step size and obstacle bypass capacity.

In MukBEF, the coiled coils fold back at the elbow position toward the SMC ATPase head, with the hinge connecting directly to one of the two heads ^6^. While ecJET-I and nvJET-II also contain an elbow, its position is clearly different, implying independent origins of these elbows. The head-to-elbow distance remains similar in ecJET-I and nvJET-II systems (≃160 Å and ≃180 Å, respectively) while being longer in MukBEF (≃286 Å) (Figure 2C). On the contrary, the elbow-to-hinge distance is similar in nvJET-II and MukBEF, but shorter in ecJET-I. In nvJET-II, the coiled coils thus extend beyond and position the hinge distant from the ATPase heads, preventing direct hinge-head contact. The hinge-proximal coiled coils contact both the heads and the KITE dimer (Figure 1B, 1C). This sandwiching may stabilize nvJET-II complexes through a more compact resting state architecture, possibly playing a role in regulating plasmid restriction.

While the Wadjet-I hinge is predicted to contain a central channel similar to many other hinges (*e.g.*, Smc5/6, Figure 2B, S5B), the nvJET-II hinge is more compact and resembles the rectangular-shaped hinge of MukBEF ^6^. The nuclease subunits also share a similar dimer architecture as predicted by AlphaFold3 (Figure S6B). Despite differences in ion requirements and likely variations in how they connect to the SMC motor units (Figure S6C, Figure 4A), the nucleases appear to serve equivalent functions in both type of Wadjet systems.

### Evolution of prokaryotic SMCs

Wadjet systems presumably undergo rapid evolutionary diversification due to ongoing biological conflicts, with selection pressures exerted by plasmid-borne anti-Wadjet counter defense mechanisms, and due to their otherwise non-essential nature. Consequently, these systems may effectively serve as natural laboratories for SMC motor evolution, where innovations emerging in one Wadjet system could be adopted by and evolve into other SMCs. While most prokaryotes rely on Smc-ScpAB for chromosome organization, certain γ-proteobacterial lineages utilize MukBEF instead ^19^. The evolutionary origin of this transition is unclear. One hypothesis suggests that MukBEF, and likely also MksBEF, originated from the domestication of Wadjet systems for host genome maintenance ^1,22–25^. This adaptation might require minimal specific modifications, as Wadjet systems likely contribute to chromosome segregation by performing DNA loop extrusion also on the host chromosome. The loss of the nuclease subunit may lock the former Wadjet system as chromosome segregation module.

Prokaryotic SMC complexes generally contain KITE homodimers, eukaryotic Smc5/6 complexes contain a KITE heterodimer (Nse1/3) ^3^, while Wadjet-II harbour a tandem-KITE, suggesting KITE diversification during evolution. The fusion of subunits followed by sequence diversification has been proposed to facilitate evolutionary innovation, as suggested for histone protein evolution ^38^. MukBEF is inhibited by the dsDNA-mimic Gp5.9 from bacteriophage T7, which targets the MukE KITE subunit ^25^. Tandem KITE proteins may have evolved under pressure from such a counter defence mechanism, facilitating escape by increasing the sequence space through KITE gene duplication. Which other Wadjet subunits experience selection pressures from anti-Wadjet mechanisms remains an intriguing question for future investigation. The deviant hinge structure of nvJET-II and the multitude of variations in the coiled coils—including in the lengths and elbow positioning— indicate that they are potentially targeted by anti-Wadjet mechanisms.

## Acknowledgments

The cryo-EM grid preparation, data collection, and initial image processing was performed at the Dubochet Center for Imaging (DCI) in Lausanne (a common initiative from EPFL, UNIGE, and UNIL) with the help of Alexander Myasnikov, Bertrand Beckert, Sergey Nazarov, Inayathulla Mohammed and Emiko Uchikawa. We are also grateful to the DCI for support during subsequent data processing, and to Kyle Muir for additional feedback and suggestions. We thank Emmanuel Jeanvoine (UNIL DCSR, *Division de Calcul et Soutien à la Recherche*) for IT support, and all members of the Gruber lab for feedback during manuscript preparation. This work was supported by the European Research Council (724482 to S.G.) and an SNSF project grant (10001017 to S.G.). F.R.-H. and H.W.L. were supported by EMBO Postdoctoral fellowships (ATLF 302-2022 and ALTF 490-2021).

## Author contributions

F.R.-H. purified the nvJetABCD proteins, performed the biochemical characterisation, carried out the cryo-EM data analysis and the model building. H.W.L. performed the *in vivo* chromosome excision assay and characterized the nature of Wadjet-II cleaved DNA ends. F.R.-H. and S.G. wrote the initial draft, and F.R.-H., H.W.L. and S.G. revised the manuscript. S.G. acquired funding and supervised the project.

## Methods

### Protein purification and protein complex reconstitution

#### Purification of nvJetABCD

JetABC and JetD from *Neobacillus vireti* LMG 21834 were produced separately in *E. coli* BL21. nvJetABC was expressed from a single co-expression vector (Table S2) with JetA amino-terminally tagged using a 10His-TwinStrep-3C affinity tag. nvJetD was expressed with a carboxy-terminal 3C-TwinStrep-10His-tag (Table S2). The cells were grown in one litre of TB-medium at 37°C until the culture reached OD_600_=0.5. The cultures were then cooled down to 16°C and protein production was induced by IPTG addition (0.25 mM final) for 16 hours. Cells were harvested by centrifugation and resuspended in lysis buffer (Tris-HCl pH 7.5 50 mM, NaCl 300 mM, glycerol 5 % (v/v), imidazole 25 mM) freshly supplemented with PMSF (1 mM) and β-mercaptoethanol (5 mM). Cells were lysed by sonication on ice with a VS70T tip using a SonoPuls unit (Bandelin), at 40 % output for 15 min with pulsing (1 sec on / 1 sec off). The lysate was then clarified by ultracentrifugation (40,000 g for 30 min). For nvJetABC, the clarified lysate was loaded onto a 5 mL StrepTrapXT affinity column, followed by 5 column volumes (CV) of washing with lysis buffer (50 mM Tris–HCl pH 7.5, 300 mM NaCl, 5 % (v/v) glycerol, 25 mM imidazole). The complex was eluted in 1 mL fractions with 4 CV of elution buffer (20 mM Tris–HCl pH 8, 200 mM NaCl, 50 mM biotin). Best fractions were pooled, and the tag was removed by addition of 3C protease (200 µL at 1 mg/mL for about ∼7mL pooled protein fractions) followed by an overnight incubation at 4°C. The resulting solution was concentrated using Amicon Ultracentrifugal filter units (50 kDa cutoff) and injected onto a Superose6 Increase 10/300 GL size-exclusion chromatography (SEC) column equilibrated with 20 mM Tris–HCl pH 7.5, 250 mM NaCl and 1 mM TCEP. Resulting fractions were concentrated to around 7.5-10 μM JetABC dimer and flash-frozen in liquid nitrogen. For nvJetD, the clarified lysate was loaded onto a 5 mL HisTrap column (Cytiva) followed by a 10 CV wash in lysis buffer. Gradient elution (10 CV) was performed with lysis buffer supplemented with 300 mM imidazole. Fractions corresponding to nvJetD were pooled and 3C protease was added to cleave the tag. At the same time, the samples were dialyzed overnight at 4°C into 20 mM Tris–HCl pH 7.5, 100 mM NaCl and 5 mM β-mercaptoethanol. Then, the protein was loaded onto a 5 mL HiTrap Q HP (Cytiva). After a 5 CV wash with fresh dialysis buffer, a gradient elution was performed using elution buffer (20 mM Tris pH 7.5, 1000 mM NaCl and 5 mM β-mercaptoethanol). Fractions containing nvJetD were then concentrated and injected onto a HiLoad Superdex200 in buffer 20 mM Tris–HCl pH 7.5, 250 mM NaCl and 1 mM TCEP. Relevant fractions were concentrated, and flash frozen in liquid nitrogen.

#### Purification of Mre11-Rad50

Mre11-Rad50 was purified as described in ^28^ (Table S2).

#### Reconstitution of nvJetABCD

nvJetABCD was reconstituted as previously described for ecJetABCD in ^5,28^ by mixing nvJetABC and nvJetD in ATG buffer (10 mM Hepes-KOH pH 7.5, 150 mM KOAc, 5 mM MgCl2, and 1 mM TCEP) supplemented with MnCl2 (5 mM or the indicated concentration) at an appropriate concentration (typically 750 nM of nvJetABC dimer with 1500 nM of nvJetD).

### DNA substrate preparation

#### Artificial DNA circles

The 2.3 kb end-biotinylated DNA substrates preparation and artificial circle formation by streptavidin were performed as in ^28^, except that the ATG buffer was supplemented with 5 mM MnCl2 prior to streptavidin addition.

### DNA cleavage assay

Wadjet cleavage reactions were performed similarly as in ^5^. Unless stated otherwise, DNA (1.8 kb at 8.75 nM) was incubated with nvJetABC (25 nM dimer) or ecJetABC (12.5 nM dimer) and nvJetD (50 nM dimer) or nvJetABC (25 nM dimer) in ATG-Mn buffer (10 mM Hepes-KOH pH 7.5, 150 mM KoAc, 5 mM MgCl_2_, 5 mM MnCl_2_), or ATG with indicated quantity of MnCl_2_, supplemented with 1 mM ATP and 1 mM TCEP in 15-30 µL reactions. The reaction was incubated at 37°C for 10 min, then arrested by addition of loading dye containing SDS (Thermofisher) and incubated for 5 min at 70°C. For reaction involving restriction enzymes, 5-10 U (BsaI-HF or RecBCD) were typically used under the same reaction conditions. For experiments involving artificial DNA circles, the DNA was subjected to Wadjet or ecMR (Mre11/Rad50, 125 nM tetramer) under the same reaction conditions. In the latter case, a loading dye without SDS was used and the sample was not incubated at 70°C to preserve streptavidin-DNA interactions. The resulting DNA species were resolved in 1% agarose gels (without SDS) containing EtBr and imaged on a transilluminator (U:GENIUS3 with a Syngene CAM-FLXCM-1 camera). The plasmids used in this study are listed in (Table S2).

For cleavage product DNA end analysis, ecJET-I, nvJET-II (alongside ScaI-HF as a blunt-end cutter control) cleaved pDonor DNA were first column purified. 600 ng of DNA were blunted by incubation with Phusion polymerase (0.5 U) for 5 minutes at 72°C (Phusion polymerase buffer, 20 µL reactions) followed by column purification. 100 ng of purified DNA was then ligated into 15 ng of a linear pJET1.2 vector (ampicillin resistant) (obtained by EcoRV-HF digestion of circular pJET1.2 followed by column purification) for 5 minutes at 22°C with T4 DNA ligase (5 U) and transformed into chemically competent *E. coli* DH5α. Plasmids were extracted from ampicillin resistant clones and subject to Sanger sequencing with primers flanking the EcoRV site (insert-vector junction). 17 clones containing full-length inserts (i.e. from a single cleavage event by nvJET-II) were analyzed. Cleavage sites were inferred upon inspection of the sequence at the junction, as well as fill-in events (indicating 5’ overhang cleavage events) (Figure S7, Table S1).

### ATPase assay

The ATP hydrolysis activity of nvJetABCD (and ecJetABCD, as control) was measured by a pyruvate kinase/lactate dehydrogenase assay at 37°C for 1 h in ATG buffer (ecJetABC/D) or ATG-Mn buffer (nvJetABC/D) (both lacking TCEP) as in Liu et al., 2022. The final concentration of JetABC (dimer) and JetD (dimer) were 62.5 nM and 125 nM, respectively. The DNA (1.8 kb, circular) was added at 33.3 nM final. The reaction was monitored by measuring the absorbance evolution at 340 nm every minute using a Synergy Neo Hybrid Multi-Mode Microplate reader, and the results were analysed using Excel and GraphPad Prism V10.2.3.

### *in vivo* chromosome excision assay

Chromosome excision assays were performed as described in ^5^. See Table S3 for the strains used.

### Cryo-EM analysis

#### Grid preparation and data collection

Prior to grid preparation, an aliquot of nvJetABC protein was freshly injected onto a Superose 6 Increase 3.2/300 equilibrated in ATG buffer (10 mM Hepes-KOH pH 7.5, 150 mM KoAc, 5 mM MgCl2, and 1 mM TCEP). 20 μL of the peak fraction (at 1.07 μM nvJetABC dimer) was then supplemented with ATP and β-octyl glucoside (final concentration of 1 mM and 0.05 % (w/v), respectively), incubated first on ice for 30 min and then at room temperature for 10 min. Cryo-EM grids (Au-flat 1.2/1.3 on 300 gold mesh) were freshly glow discharged using an EasyGlow device with 15 mA current, 90 s glow time and 60 s wait time. Next, 3 µl of the nvJetABC sample was applied onto the grids mounted in a Vitrobot Mark IV, with a chamber set to 10°C and 100% humidity. The grids were blotted at blot force 10 for 0.5 s and vitrified in liquid ethane precooled to liquid nitrogen temperature. Two grids were made from the same sample. Grids screening followed by exploratory dataset acquisition (3,528 movies collected, EER format) were performed on a Glacios Cryo-TEM equipped with Falcon IV CMOS detector (Thermofisher Scientific (TFS)) operated by the EPU software (TFS) at a magnification of 150,000x (nominal pixel of 0.92 Å) with a total dose of 40 electrons per square angstrom (e-/Å2) and using a defocus range from -1 to -2.4 µm. High-resolution collection (from two duplicate grids) was performed on a 300 kV Titan Krios equipped with a Falcon IV G4i camera (TFS) and a SelectriX energy filter (10 eV slit) at a magnification of 165,000x (pixel size of 0.726 Å), operated by the EPU software (TFS). 22,188 movies (EER format) were collected with a total dose of 40 e-/Å2 and using a defocus range from -0.8 to -2.6 µm.

#### Data processing

Initial data processing was performed on-the-fly using cryoSPARC-live and further processing was done in cryoSPARC (initially V3.3 then upgraded to V4.5) (Figure S2, S3, S4, Table 1). 22,188 dose-weighted micrographs were imported in cryoSPARC and after CTF estimation, 19,566 micrographs were kept based on ice quality and CTF fit resolution. Particle picking was performed using the blob picker (elliptical) and led to the picking of 10,792,128 initial particles that were extracted with a box size of 600 pixels (Fourier-cropped 8 times; 7,960,948 extracted particles). A final stack of 385,630 particles were selected after several rounds of 2D classification and particle re-extraction with a box of 600 pixels (Fourier-cropped 2 times). We found that particle re-picking using the template picker did not significantly improve the particle stack. The 385,630 particles were subjected to *ab initio* reconstruction with 3 classes. After duplicate removal, non-uniform refinement of the best class gave an initial reconstruction of a single nvJetABC motor unit at 3.68 Å (151,158 particles). 3D variability analysis suggested significant flexible motion within the complex, especially in the nvJetC coiled coils and in kleisin. Thus, we performed 3D classification (5 classes) with this set of particles. The best resolved class contained 47,279 particles that were subjected to a non-uniform refinement, followed by particles re-extraction with centring (box size: 640 pixels, Fourier-cropped to 400 pixels). A final non-uniform refinement with global CTF correction gave the 3D reconstruction of a single nvJetABC motor unit at 3.5 Å. While the map represents a single SMC motor unit, fuzzy density belonging to the second motor unit (note that the nvJetA kleisin nWHD from the second SMC motor unit is visible and included in the deposited model) was discernible close to the kleisin N-terminus. Such fuzzy density was also clearly visible in several 2D classes (Figure S2). To determine a structure of the holo-complex, the particle set of the initial reconstruction (151,158 particles) was first re-extracted with a larger box size of 1,000 pixel (Fourier-cropped 2 times) and subjected to 2D classification in order to identify particles that best align as a dimer of SMC motor units. The resulting 68,967 particles were subjected to a double class *ab initio* reconstruction. The best class contained improved density in the region belonging to the second motor. Then, the particles set belonging to the initial reconstruction (151,158 particles) were extracted at a box size of 800 (Fourier crop 2 times) and aligned on the previously improved *ab initio* reference. A non-uniform refinement with C2 symmetry enforced gave a poorly resolved reconstruction of the full complex, that was used to create an extended mask containing both SMC motor units. The particle set (aligned to in C1 by a non-uniform refinement) was then subjected to 3D classification (5 classes) using the previously generated mask containing both motor units of the holocomplex as a solvent mask. The best resolved class (15,908 particles) was subjected to non-uniform refinement with C1 symmetry. The reconstruction clearly showed density for both SMC motor units (albeit one of them was less well resolved). A final non-uniform refinement with C2 symmetry enforced gave the 3D reconstruction of the full nvJetABC holocomplex (dimer) at an estimated overall resolution of 6.71 Å. Both C1 and C2-enforced maps were found to overlap well, with the second motor unit much better resolved in the C2 map. To obtain the local map of the nvJetC hinge, the 151,158 particles from the initial 3D reconstruction were subjected to a local classification (5 classes) with a mask solely including the hinge region and a solvent mask around the whole SMC motor unit. The best resolved class (34,013 particles) was subjected to local refinement, which gave a 3D reconstruction of the nvJetC hinge at low resolution (estimated overall resolution: 4.38 Å). All reconstruction were subjected to local resolution estimation in cryoSPARC (Figure S3).

#### Model building

To build the model of the nvJetABC monomer, AlphaFold2 predictions run on the UNIL computing cluster or on Colabfold ^39^ (isolated parts were predicted, *e.g.* the upper part of the nvJetC coiled coils – the hinge, nvJetC head – the lower part of coiled coils, nvJetA-nvJetB, nvJetA Nt-nvJetC-head) were used as starting material. The AlphaFold2 models were segmented, rigid body docked, flexibly fitted and rebuilt into the nvJetABC cryo-EM density using ChimeraX (V1.4 then updated to V1.8) ^40^, ISOLDE (V1.6 then updated to V1.8) ^41^ and Coot ^42^. The model was iteratively improved using ISOLDE/Coot and real space refinement in PHENIX (V1.20.1) ^43,44^. Finally, the model was validated using MolProbity ^45^ and EMringer ^46^ implementations in PHENIX. Figure preparation was done using ChimeraX and Chimera ^47^. 3D FSC sphericity values were obtained with the 3DFSC^48^ program run on the remote 3DFSC processing server (New York Structural Biology Center, Salk Institute for Biological Studies). The Q-score reported in Table 1 ^49^ was obtained from wwPDB during model deposition. For the model of nvJetABC dimer, the model of nvJetABC monomer was first flexibly fitted into the dimer map using ISOLDE under distances and secondary structures restraints. After manual correction using ISOLDE, the model was subjected to real space refinement in PHENIX. The aim of this model is to illustrate the overall geometry of the nvJetABC dimer. Given the lower resolution of the corresponding map, we suggest the reader to refer to the JetABC monomer model (PDB: 9QE1) for any detailed analysis instead.

