## Supplemental Figures and Legends for "Structure of a Type II SMC Wadjet Complex from *Neobacillus vireti*": Supplemental_figures_and_legends.pdf

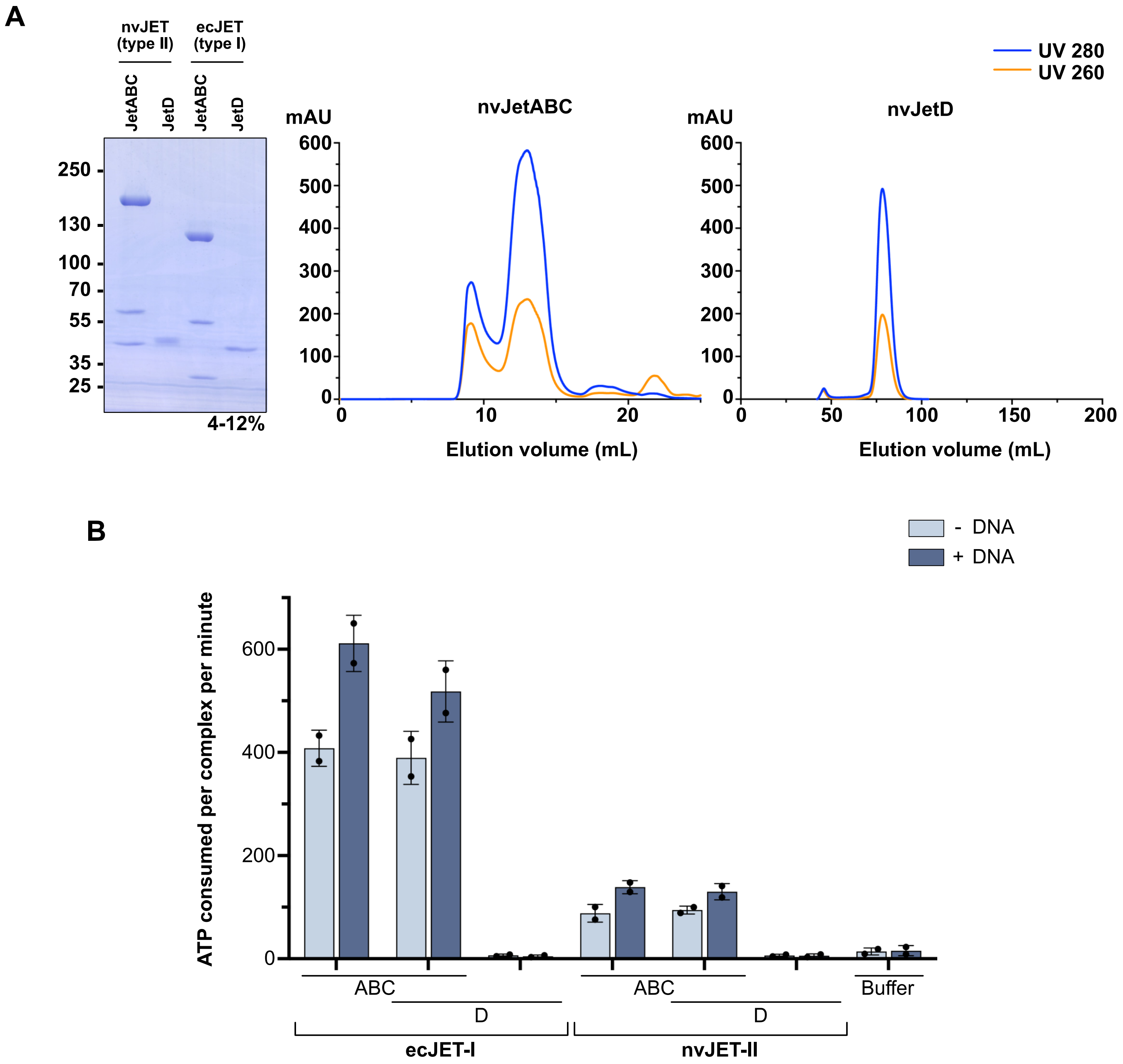
A
nvJET (type II)
ecJET (type I)
JetABC
JetD
JetABC
JetD
250
130
100
70
55
35
25
4-12%
mAU
nvJetABC
600
500
400
300
200
100
0
0
10
20
Elution volume (mL)
mAU
nvJetD
0
50
100
150
200
Elution volume (mL)
UV 280
UV 260
B
- DNA
+ DNA
ATP consumed per complex per minute
600
400
200
0
ABC
D
ecJET-I
ABC
D
nvJET-II
Buffer

**Figure S1**

**Figure S1: Reconstitution of a type II Wadjet system from *Neobacillus vireti*.** Related to Figure 1.

(A) Left: Migration profiles of purified nvJET-II proteins compared to ecJET-I [5]. Proteins have been diluted respectively to 1.25 µM (JetABCD) and 3.5 (JetD) µM prior to loading on a 4-12% gradient acrylamide gel (Coomassie brilliant blue stained). Right: representative examples of the size exclusion chromatography profiles of nvJetABC and nvJetD on a Superose 6 Increase 10/300 GL and HiLoadSuperdex200, respectively. (B) ATP-hydrolysis assay with JetABC from the indicated species (62.5 nM JetABC dimer, 125 nM JetD dimer, 33.3 nM pDonor plasmid DNA). The graph bars represent the mean from two independent replicates, with errors bars corresponding to the standard deviation.

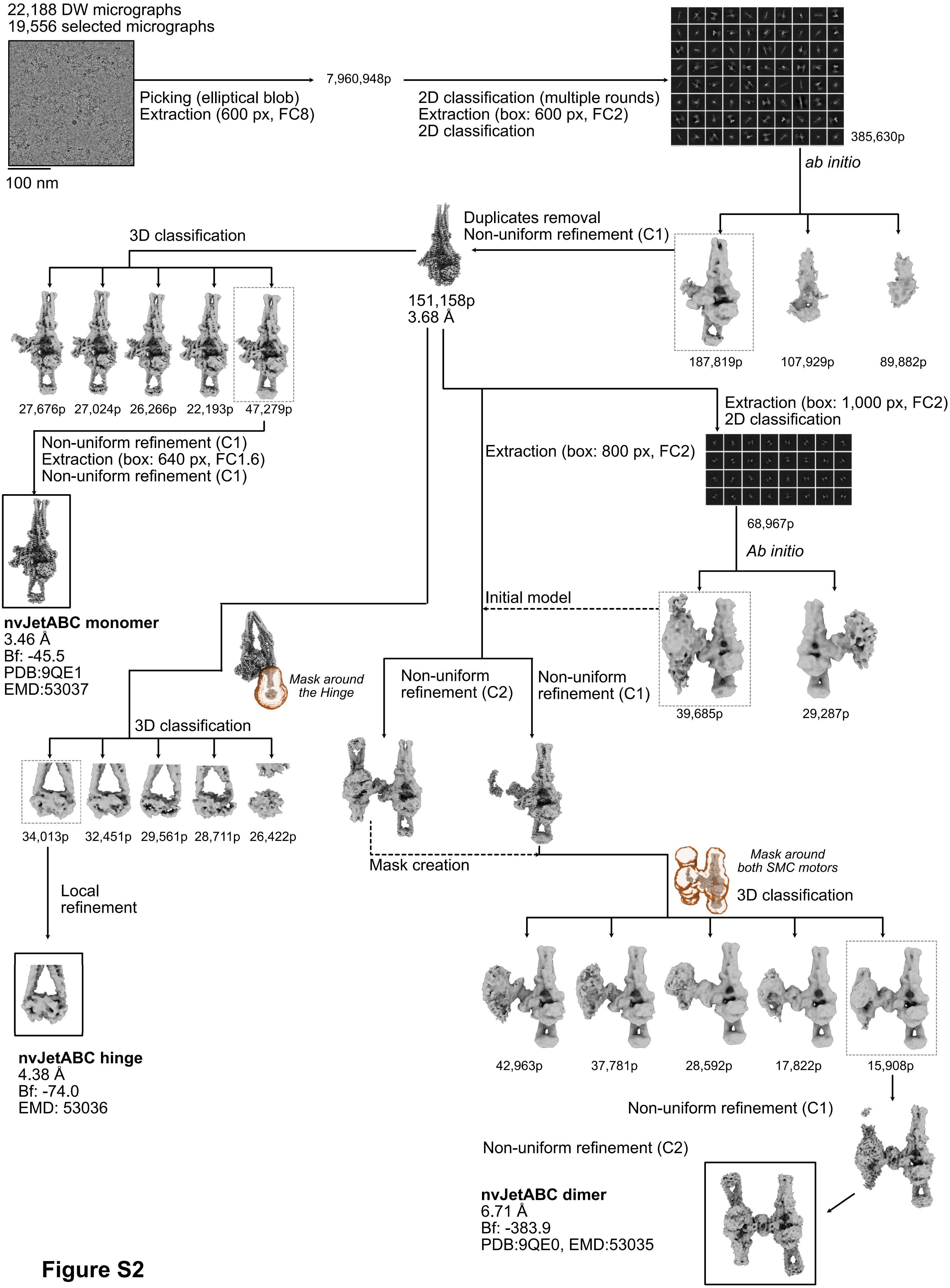

22,188 DW micrographs
19,556 selected micrographs
100 nm
Picking (elliptical blob)
Extraction (600 px, FC8)
7,960,948p
2D classification (multiple rounds)
Extraction (box: 600 px, FC2)
2D classification
385,630p
ab initio
Duplicates removal
Non-uniform refinement (C1)
187,819p
107,929p
89,882p
151,158p
3.68 Å
3D classification
27,676p
27,024p
26,266p
22,193p
47,279p
Non-uniform refinement (C1)
Extraction (box: 640 px, FC1.6)
Non-uniform refinement (C1)
nvJetABC monomer
3.46 Å
Bf: -45.5
PDB:9QE1
EMD:53037
Extraction (box: 1,000 px, FC2)
2D classification
Extraction (box: 800 px, FC2)
68,967p
Ab initio
Initial model
39,685p
29,287p
Mask around the Hinge
3D classification
Non-uniform refinement (C2)
Non-uniform refinement (C1)
34,013p
32,451p
29,561p
28,711p
26,422p
Local refinement
Mask creation
Mask around both SMC motors
3D classification
nvJetABC hinge
4.38 Å
Bf: -74.0
EMD: 53036
42,963p
37,781p
28,592p
17,822p
15,908p
Non-uniform refinement (C1)
Non-uniform refinement (C2)
nvJetABC dimer
6.71 Å
Bf: -383.9
PDB:9QE0, EMD:53035


**Figure S2**

**Figure S2: Cryo-EM analysis pipeline.** Related to Figure 1.

Representative dose-weighted micrograph (low-pass filtered to 5 Å) and 2D classes are shown. The abbreviations used in the figure are: DW: dose-weighted; Box: extraction box; px: pixels: FC: Fourier-crop; Bf: sharpening B-factor. The final maps that were obtained are shown in a black box. See Figure S3 for local resolution estimation and FSC curves, and Figure S4 for model fitting into their respective maps.

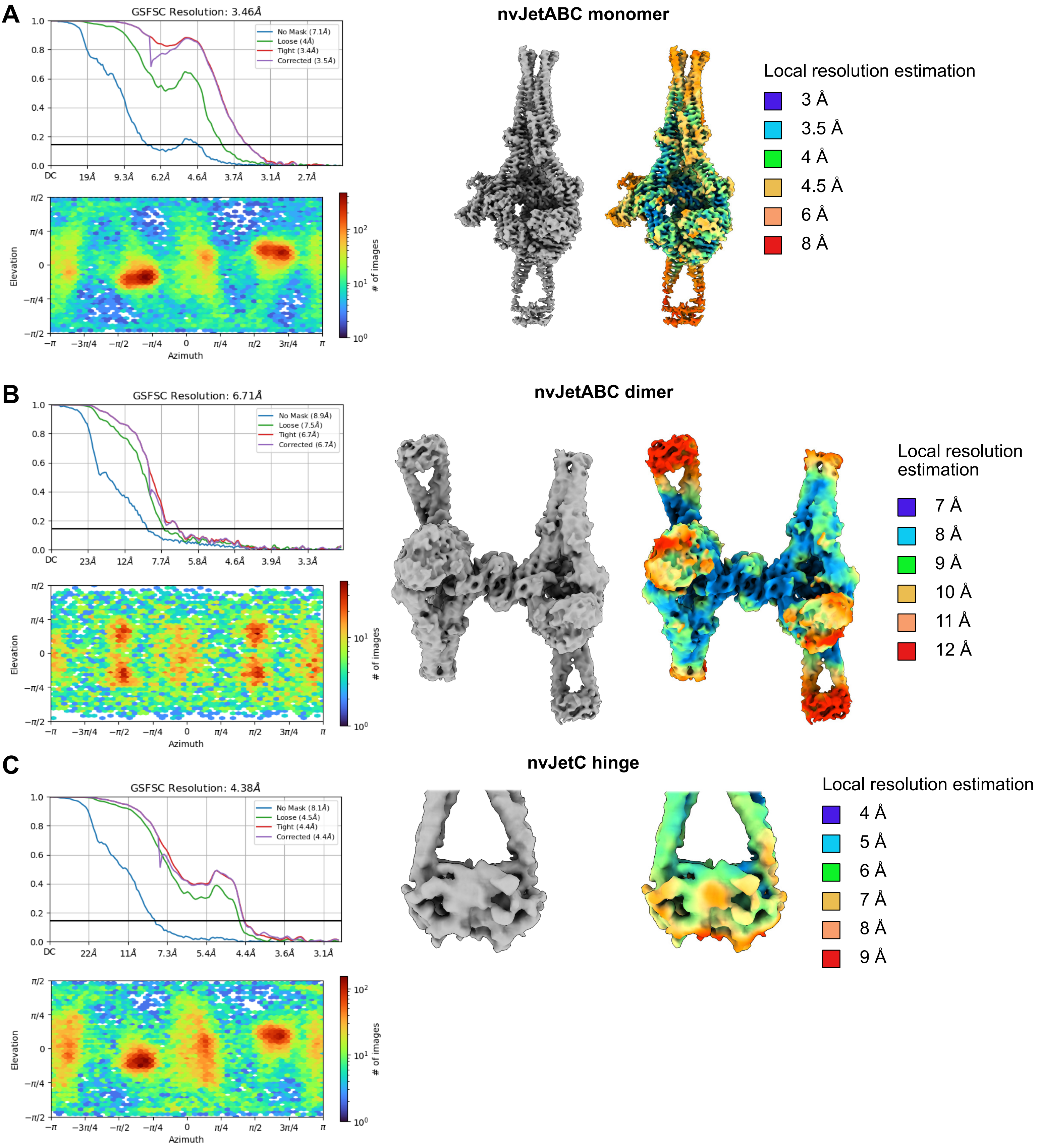
A
GSFSC Resolution: 3.46Å
No Mask (7.1Å)
Loose (4Å)
Tight (3.4Å)
Corrected (3.5Å)
Elevation
Azimuth
### of images
nvJetABC monomer
Local resolution estimation
3 Å
3.5 Å
4 Å
4.5 Å
6 Å
8 Å
B
GSFSC Resolution: 6.71Å
No Mask (8.9Å)
Loose (7.5Å)
Tight (6.7Å)
Corrected (6.7Å)
nvJetABC dimer
Local resolution estimation
7 Å
8 Å
9 Å
10 Å
11 Å
12 Å
C
GSFSC Resolution: 4.38Å
No Mask (8.1Å)
Loose (4.5Å)
Tight (4.4Å)
Corrected (4.4Å)
nvJetC hinge
Local resolution estimation
4 Å
5 Å
6 Å
7 Å
8 Å
9 Å

**Figure S3**

**Figure S3: Cryo-EM maps resolution estimation.** Related to Figure 1.

FSC curves, orientation distribution and local resolution estimation (as computed in cryoSPARC) are shown for (A) the nvJetABC monomer map, (B) the nvJetABC dimer map and (C) the nvJetC hinge map.

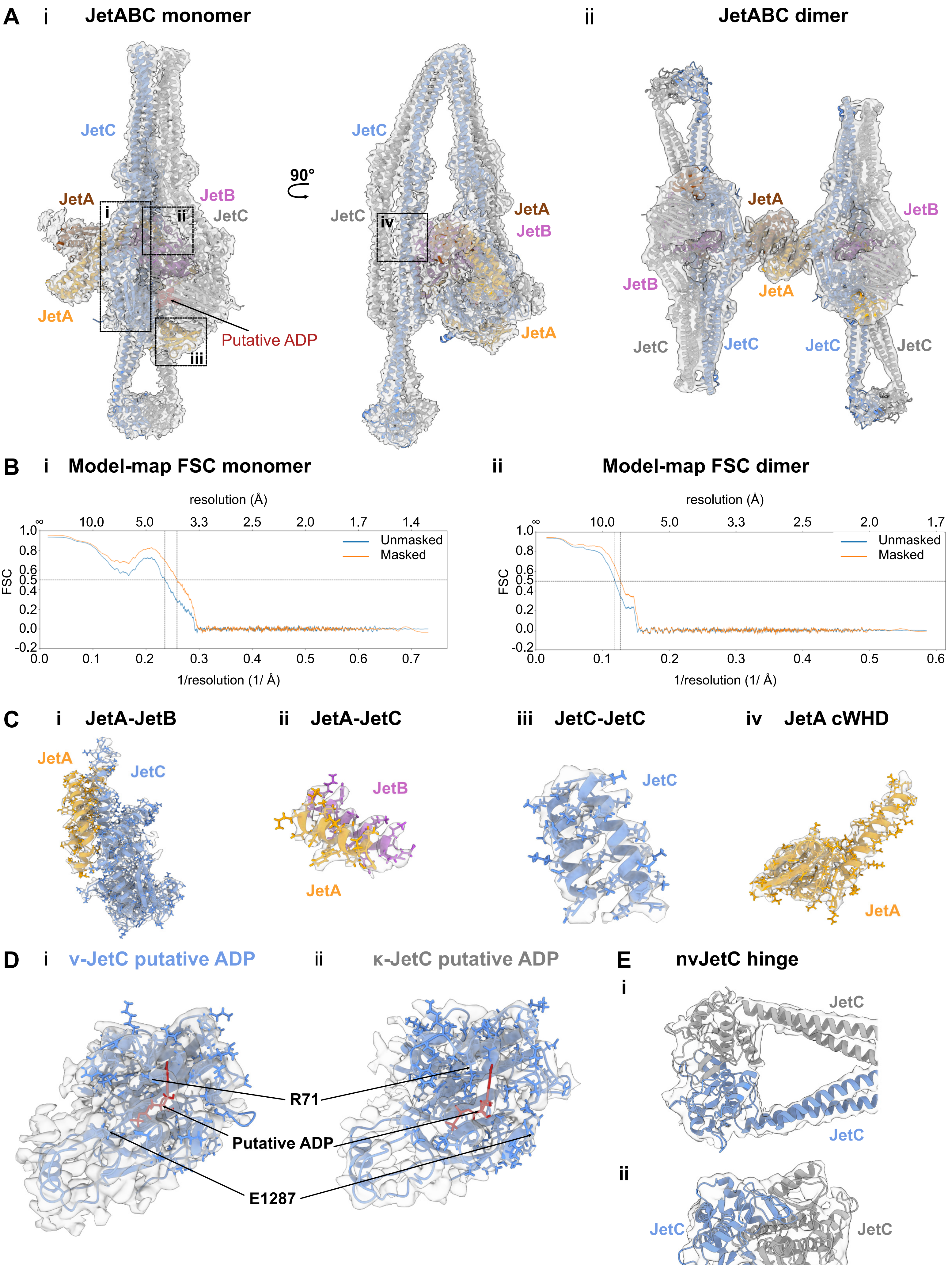

A
i
JetABC monomer
JetC
JetA
JetB
JetC
i
ii
iii
JetA
Putative ADP
90°
JetC
JetC
iv
JetA
JetB
JetA
ii
JetABC dimer
JetA
JetB
JetB
JetA
JetC
JetC
JetC
JetC
B
i
Model-map FSC monomer
resolution (Å)
∞
10.0
5.0
3.3
2.5
2.0
1.7
1.4
Unmasked
Masked
FSC
1/resolution (1/ Å)
ii
Model-map FSC dimer
resolution (Å)
10.0
5.0
3.3
2.5
2.0
1.7
Unmasked
Masked
FSC
1/resolution (1/ Å)
C
i
JetA-JetB
JetA
JetC
ii
JetA-JetC
JetB
JetA
iii
JetC-JetC
JetC
iv
JetA cWHD
JetA
D
i
v-JetC putative ADP
ii
κ-JetC putative ADP
R71
Putative ADP
E1287
E
nvJetC hinge
i
JetC
JetC
ii
JetC
JetC


**Figure S4**

**Figure S4: Models fit to map.** Related to Figure 1.

(A) Overall fit of the models into their respective cryo-EM densities for (i) the nvJetABC monomer (PDB-9QE1) and (ii) the nvJetABC dimer (PDB-9QE0). (B) Model-map FSC curves as computed during the final refinement in PHENIX for (i) the nvJetABC monomer and (ii) the nvJetABC dimer. (C) Selected regions of (A) showing the model fitting into the cryo-EM map of the nvJetABC monomer (PDB: 9QE1). (D) Cryo-EM density corresponding to a nucleotide in the two nvJetC subunits (monomer, PDB: 9QE1). (E) AlphaFold2 model of nvJET-II hinge rigid body fitted into its cryo-EM map (EMD-53036). (i) View from the side, (ii) view from the top.

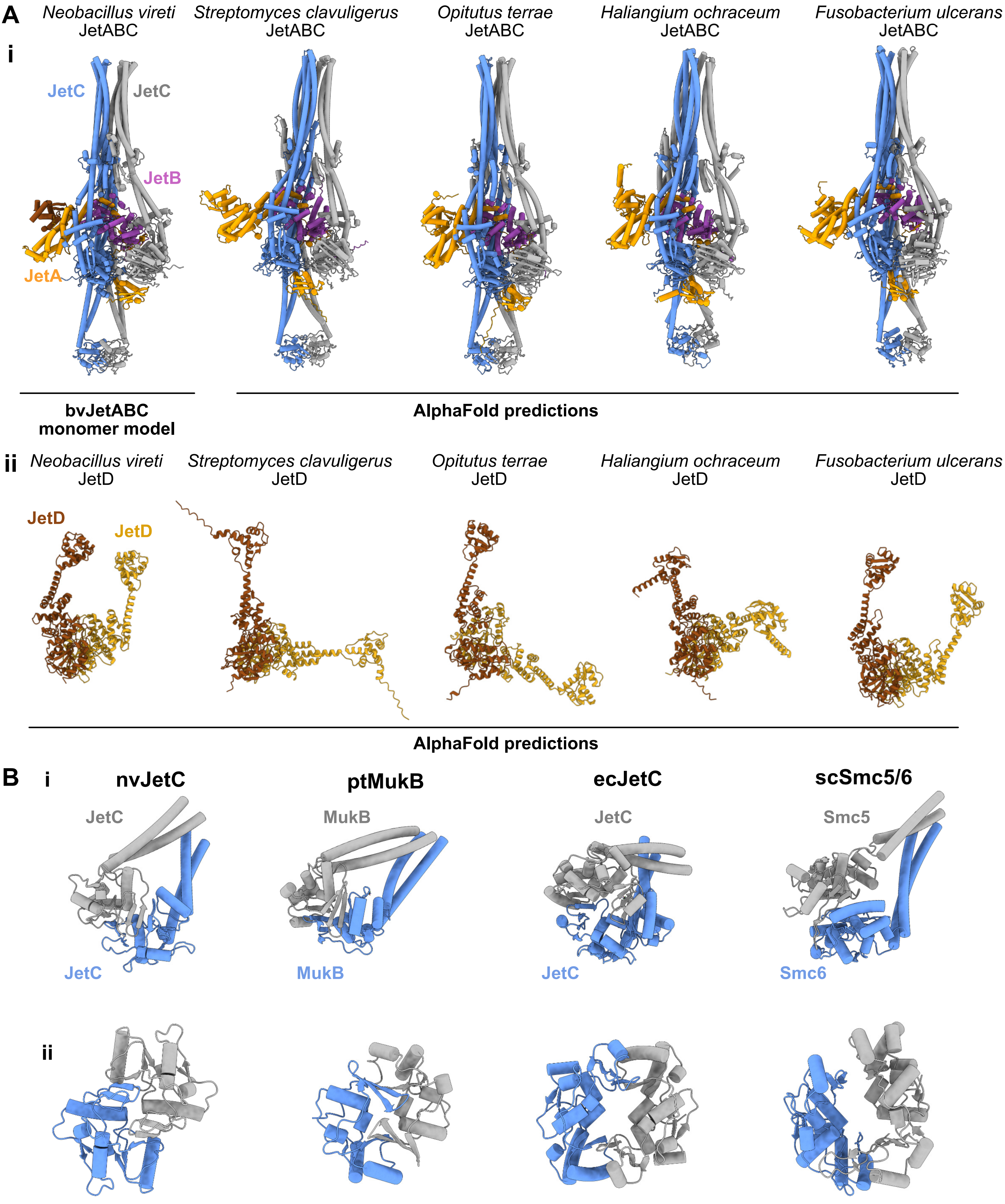
A
i
Neobacillus vireti
JetABC
Streptomyces clavuligerus
JetABC
Opitutus terrae
JetABC
Haliangium ochraceum
JetABC
Fusobacterium ulcerans
JetABC
JetC
JetC
JetB
JetA
bvJetABC
monomer model
AlphaFold predictions
ii
Neobacillus vireti
JetD
Streptomyces clavuligerus
JetD
Opitutus terrae
JetD
Haliangium ochraceum
JetD
Fusobacterium ulcerans
JetD
JetD
JetD
AlphaFold predictions
B
i
nvJetC
ptMukB
ecJetC
scSmc5/6
JetC
MukB
JetC
Smc5
JetC
MukB
JetC
Smc6
ii

**Figure S5**

**Figure S5: AlphaFold predictions of Wadjet-II systems.** Related to Figure 2.

(A) AlphaFold3 predictions of selected Wadjet-II systems (*Streptomyces clavuligerus*, *Opitutus terrae*, *Haliangium ochraceum*, *Fusobacterium ulcerans*) from [26]. (i) JetABC SMC motor units; (ii) JetD nuclease dimers. The selected AlphaFold3 predictions are compared with the nvJetABC monomer model (PDB: 9QE1) and the nvJetD AlphaFold2 model. (B) Comparison of hinge shapes (and lumens) of nvJetC, ecJetC (AlphaFold2 predictions), ptMukB (PDB: 7NZ4) and scSmc5/6 (PDB: 7QCD). The hinge domain is displayed, with SMC monomers labelled in blue and grey colors, respectively. (i) Display of the hinge domain with the adjacent coiled coils portion, (ii) display centered on the lumen of the hinge.

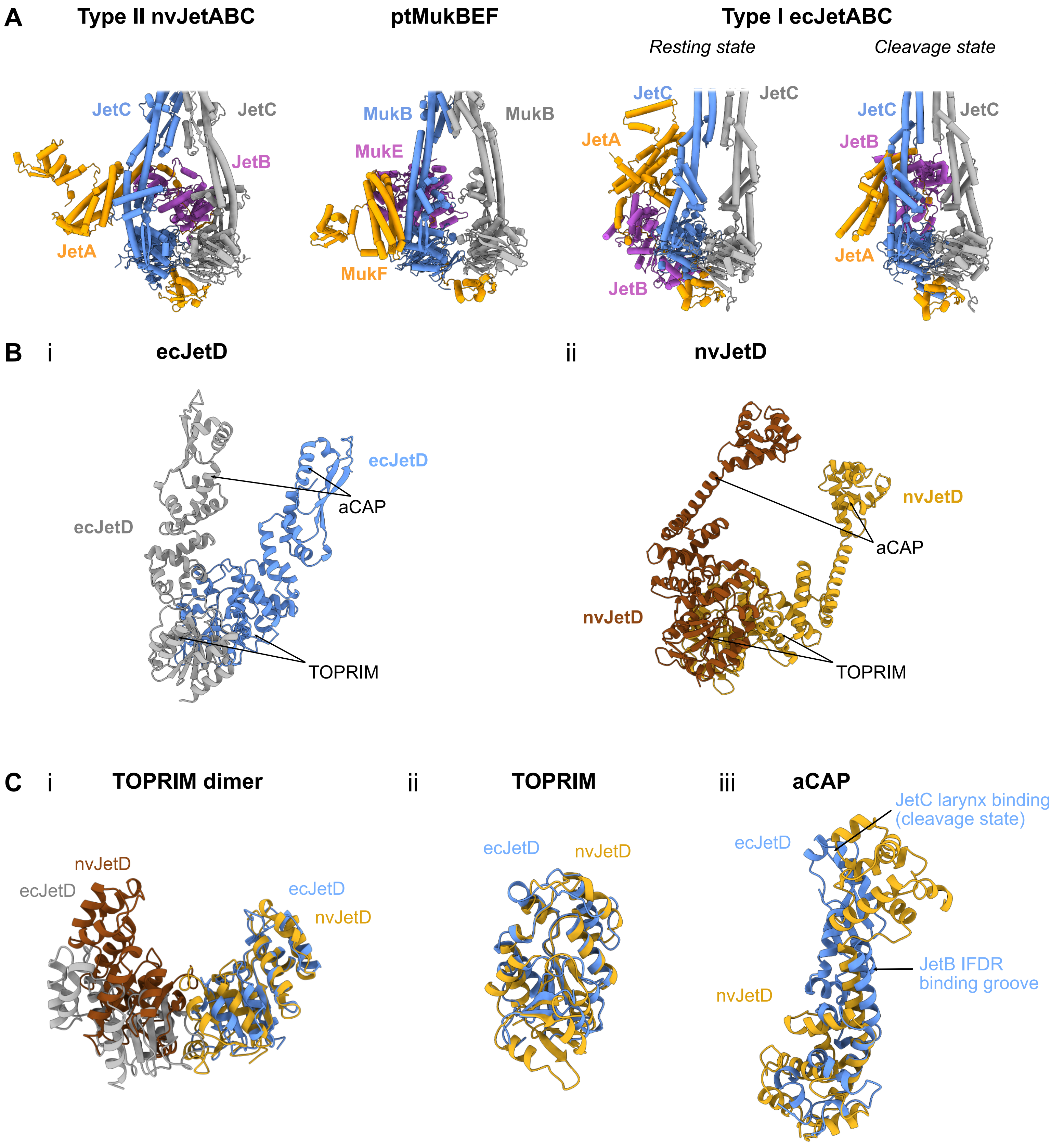
A
Type II nvJetABC
ptMukBEF
Type I ecJetABC
Resting state
Cleavage state
JetC
JetC
JetB
JetA
MukB
MukB
MukE
MukF
JetC
JetC
JetA
JetB
JetC
JetC
JetB
JetA
B
i
ecJetD
ecJetD
aCAP
ecJetD
TOPRIM
ii
nvJetD
nvJetD
aCAP
nvJetD
TOPRIM
C
i
TOPRIM dimer
nvJetD
ecJetD
ecJetD
nvJetD
ii
TOPRIM
ecJetD
nvJetD
iii
aCAP
JetC larynx binding
(cleavage state)
ecJetD
JetB IFDR
binding groove
nvJetD

**Figure S6**

**Figure S6: Structural comparison of SMC motor units and JetD dimers.** Related to Figure 2.

(A) Comparison of the organisation of SMC motor units (focusing on the head-proximal region) for nvJetABC monomer (PDB: 9QE0), ptMukBEF (PDB: 7NZ4) and ecJET-I in its resting and cleavage competent states (PDB: 8AS8 and 8Q72, respectively). (B) AlphaFold2 predictions of JetD dimers from (i) ecJET-I [5] and (ii) nvJET-II (this study). (C) Comparison of nvJetD with ecJetD (AlphaFold2 predictions). (i) superimposition of TOPRIM dimers, (ii) superimposition of an individual toprim domain, (iii) superimposition of the aCAP domains. In Wadjet-I systems, the aCAP (CAP: catabolite activator protein; aCAP: arm-CAP) domain binds to JetABC through interactions with the JetB N-terminal IFDR motif (in *P. aeruginosa* PA14 Wadjet) [4] and by inserting into the JetC larynx in the cleavage-competent state (in ecJET-I) [28].

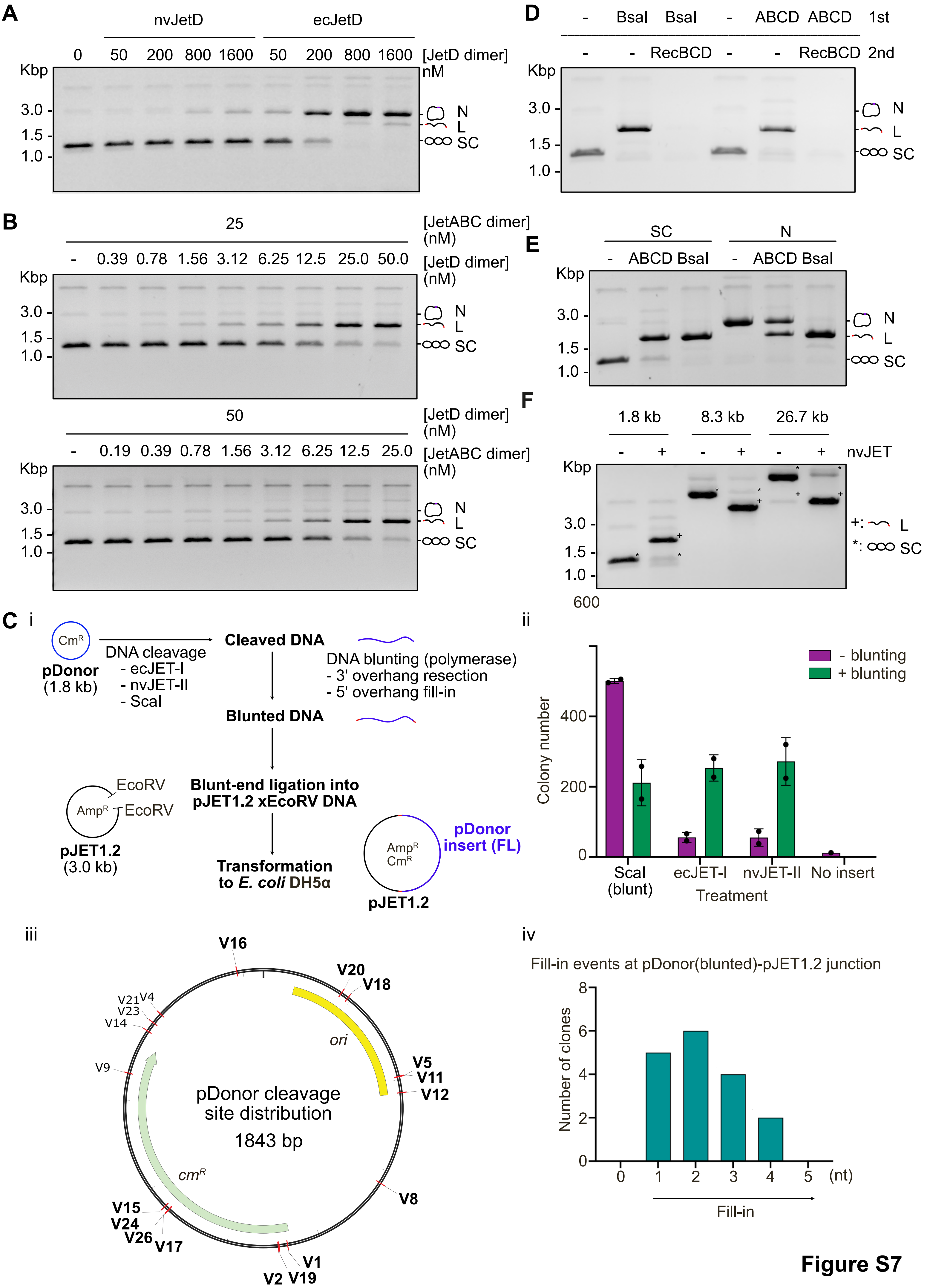
A
nvJetD
ecJetD
0 50 200 800 1600 50 200 800 1600 [JetD dimer] nM
Kbp
3.0
1.5
1.0
N
L
SC
B
25 [JetABC dimer] (nM)
- 0.39 0.78 1.56 3.12 6.25 12.5 25.0 50.0 [JetD dimer] (nM)
50 [JetD dimer] (nM)
- 0.19 0.39 0.78 1.56 3.12 6.25 12.5 25.0 [JetABC dimer] (nM)
C
i
pDonor (1.8 kb)
DNA cleavage
- ecJET-I
- nvJET-II
- ScaI
Cleaved DNA
DNA blunting (polymerase)
- 3' overhang resection
- 5' overhang fill-in
Blunted DNA
EcoRV
EcoRV
pJET1.2 (3.0 kb)
Blunt-end ligation into pJET1.2 xEcoRV DNA
Transformation to E. coli DH5α
pDonor insert (FL)
pJET1.2
ii
Colony number
- blunting
+ blunting
ScaI (blunt)
ecJET-I
nvJET-II
No insert
Treatment
iii
pDonor cleavage site distribution
1843 bp
ori
cmR
V16 V20 V18 V5 V11 V12 V8 V1 V2 V19 V15 V24 V26 V17 V9 V14 V23 V21 V4
iv
Fill-in events at pDonor(blunted)-pJET1.2 junction
Number of clones
Fill-in
(nt)
D
- BsaI BsaI - ABCD ABCD 1st
- - RecBCD - - RecBCD 2nd
E
SC
N
- ABCD BsaI - ABCD BsaI
F
1.8 kb 8.3 kb 26.7 kb
- + - + - + nvJET
+: L
*: SC
600

Figure S7

**Figure S7: Biochemical characterization of the plasmid cleavage activity of nvJET.** Related to Figure 4.

(A) Cleavage assay with ecJetD and nvJetD in presence of manganese to examine their potential promiscuous DNA nicking activities. Treatment of plasmid DNA (8.75 nM) with up to 1,600 nM of nvJetD (dimers) showed minimal nicking, whereas 200 nM of ecJetD dimers caused substantial plasmid nicking. Nevertheless, at the protein concentrations used in this study (Figure 4), ecJetD's non-specific nicking remained minimal, with plasmid cleavage fully dependent on both JetABC and JetD. The resulting DNA species are labelled: N for Nicked; L for Linear; SC for supercoiled. The same labelling is used in the following panels. (B) Cleavage assay with the indicated concentrations of nvJET-II proteins. (C) Cleavage site preferences of nvJET-II. (i) Schematic adapted from [4] of cleaved pDonor DNA blunting, and subsequent blunt-end ligation into pJET1.2 acceptor (see Methods). (ii) Counts of ampicillin-resistant colonies after transformation with pJET1.2 ligated with pDonor subject to the indicated treatments. Note that cleavage products from ecJET-I and nvJET-II that were not subject to blunting treatment still yielded a small number of clones, indicating both Wadjets can also infrequently catalyze blunt-end DNA cleavage. (iii) Distribution of pDonor cleavage sites (red). Only clones from clones containing full-length pDonor insert (derived from blunting treatment) were analyzed. (iv) Distribution of fill-in lengths (representing 5’ overhang cleavage events) from clones containing full-length pDonor insert (derived from blunting treatment). (D) Post-treatment by RecBCD (10 min at 37°C) of nvJET-II cleaved DNA resulted in elimination of all DNA species. (E) Plasmid cleavage assay with supercoiled (SC) or nicked plasmid DNA (N). (F) Cleavage assay with larger plasmids. Plasmids used: pDonor (1.8 kb, 8.75 nM), pHCM05 (8.3 kb, 1.94 nM) and pMT (26.7 kb, 0.61 nM). The resulting DNA species are labelled directly on the gel + for Linear and * for supercoiled. The experiments shown are a representative example of at least two independent replicates.

A

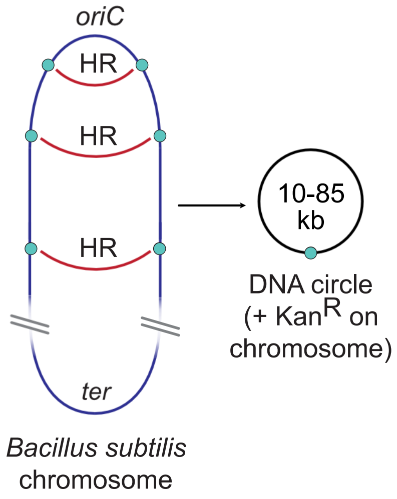
oriC
HR
HR
HR
10-85
kb
DNA circle
(+ KanR on
chromosome)
ter
Bacillus subtilis
chromosome

B

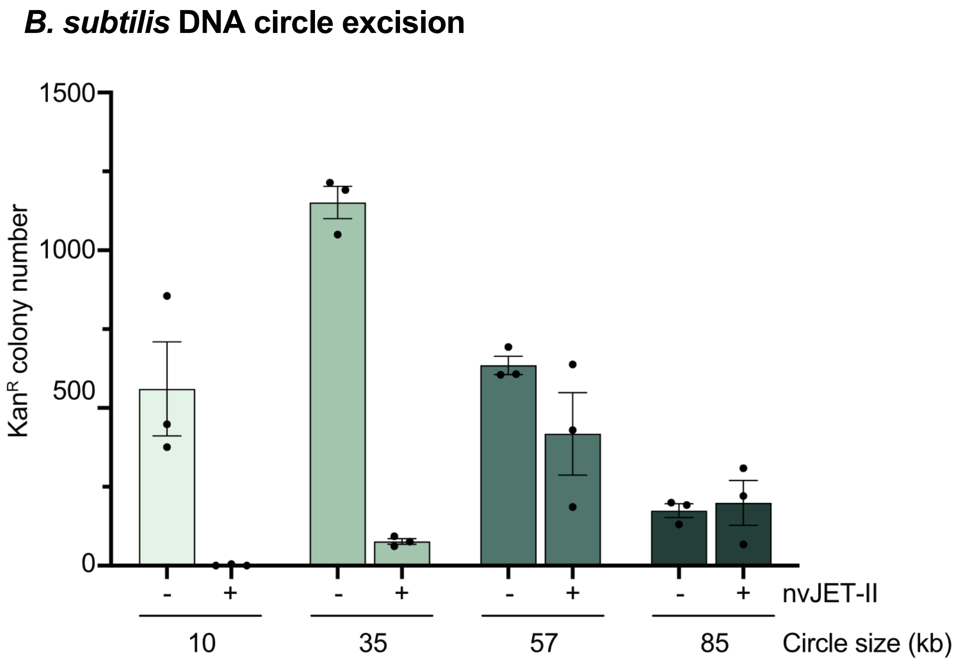
B. subtilis DNA circle excision
1500
1000
500
0
KanR colony number
- + - + - + - +
nvJET-II
10
35
57
85
Circle size (kb)

**Figure S8**

**Figure S8: Plasmid restriction size threshold of nvJET-II.** Related to Figure 4.

(A) Schematic of the chromosome excision experiment adapted from [5]. Briefly, *B. subtilis* strains harboring partially overlapping fragments of the neomycin resistance gene are placed at defined distances away from the origin of replication (*oriC*) in the same orientation. Sporadic homologous recombination events give rise to excised circles of defined size (10-85 kb). The chromosome gains a full-length neomycin resistance gene (as well as an exogenous *oriN* origin of replication), which was used as a readout for circle excision. (B) Graph showing the number of neomycin resistant colonies of the indicated strains after ~8 generations of growth. The data from three independent replicates are shown.
